# eQTL Catalogue: a compendium of uniformly processed human gene expression and splicing QTLs

**DOI:** 10.1101/2020.01.29.924266

**Authors:** Nurlan Kerimov, James D. Hayhurst, Kateryna Peikova, Jonathan R. Manning, Peter Walter, Liis Kolberg, Marija Samoviča, Manoj Pandian Sakthivel, Ivan Kuzmin, Stephen J. Trevanion, Tony Burdett, Simon Jupp, Helen Parkinson, Irene Papatheodorou, Andrew Yates, Daniel R. Zerbino, Kaur Alasoo

**Affiliations:** Institute of Computer Science, University of Tartu, Tartu, 51009, Estonia; Open Targets, South Building, Wellcome Genome Campus, Hinxton, Cambridge CB10 1SD, UK; European Molecular Biology Laboratory, European Bioinformatics Institute, Wellcome Genome Campus, Hinxton, Cambridge CB10 1SD, UK

**Author notes:** These authors contributed equally to this work (Nurlan Kerimov, James D. Hayhurst). These authors jointly supervised this work (Daniel R. Zerbino, Kaur Alasoo). Correspondence should be addressed to D.R.Z or K.A.

## Abstract

An increasing number of gene expression quantitative trait locus (eQTL) studies have made summary statistics publicly available, which can be used to gain insight into complex human traits by downstream analyses, such as fine mapping and colocalisation. However, differences between these datasets, in their variants tested, allele codings, and in the transcriptional features quantified, are a barrier to their widespread use. Consequently, target genes for most GWAS signals have still not been identified. Here, we present the eQTL Catalogue (https://www.ebi.ac.uk/eqtl/), a resource which contains quality controlled, uniformly recomputed QTLs from 21 eQTL studies. We find that for matching cell types and tissues, the eQTL effect sizes are highly reproducible between studies, enabling the integrative analysis of these data. Although most *cis*-eQTLs were shared between most bulk tissues, the analysis of purified cell types identified a greater diversity of cell-type-specific eQTLs, a subset of which also manifested as novel disease colocalisations. Our summary statistics can be downloaded by FTP, accessed via a REST API, and visualised on the Ensembl genome browser. New datasets will continuously be added to the eQTL Catalogue, enabling the systematic interpretation of human GWAS associations across many cell types and tissues.

## Introduction

Gene expression and splicing QTLs are a powerful tool to link disease-associated genetic variants to putative target genes. However, despite efforts by large-scale consortia such as GTEx *(1)* and eQTLGen *(2)* to provide comprehensive eQTL annotations for a large number of human tissues, target genes and relevant biological contexts for most GWAS signals have not been found yet. Systematic colocalisation efforts based on GTEx data have identified putative target genes for 47% of the GWAS loci *(3)*. Still, these genetic effects mediate only 11% of disease heritability *(4),* suggesting that many regulatory effects cannot be detected in bulk tissues at a steady-state *(5)*. In contrast, profiling specialised disease-relevant cell types such as induced pluripotent stem cells *(6)*, peripheral immune cells *(7)*, microglia *(8, 9)* or dopaminergic neurons *(10)* often identifies additional colocalisations that are missing in GTEx. While several databases have been developed to collect eQTL summary statistics from individual studies *(11*–*17)*, these efforts have relied on the heterogeneous set of files provided by the original authors. These results often contain only a small subset of significant associations or lack essential details such as effect alleles, standard errors or sample sizes, which limit the downstream colocalisation and Mendelian randomisation analyses that can be performed *(18)*.

Moreover, there is considerable technical variation between studies in sample collection, RNA sequencing, genotyping and data analysis. Thus, it is currently unclear how strongly eQTL effect sizes are influenced by technical differences in sample collection, how many eQTLs are broadly shared, and what fraction are specific to a given cell or tissue type and could thus give rise to novel disease colocalisations. While analyses based on GTEx data have generally estimated high levels of eQTL sharing between most bulk tissues *(1*,*19)*, smaller studies have often estimated much lower levels of sharing between purified cell types *(20, 21)*. However, these analyses are sensitive to how sharing is defined, which genes and variants are included in the analysis and which analytical approaches are used *(19, 22)*. Thus, it is impossible to directly compare the estimates of eQTL sharing between studies without re-analysing the individual-level data with uniform methods.

Recent methodological advances have made it feasible to fine map genetic associations to small credible sets of putative causal variants and distinguish between multiple independent genetic signals in the region *(23, 24)*. These fine mapping results can be directly used in colocalisation analysis *(25)*. They can also help avoid the many false negative colocalisations missed by approaches that assume a single causal variant in the region of interest *(18)*. However, reliable fine mapping requires precise information about in-sample linkage disequilibrium (LD) between genetic variants which is usually not available *(26, 27)*.

To overcome these limitations, we have uniformly re-processed (see Figure 1) individual-level eQTL data from 112 datasets across 21 independent studies (see Figure 2). We find that eQTL effect sizes from matched cell types or tissues are generally highly reproducible between studies. Using both eQTL sharing and matrix factorisation approaches on fine mapped eQTL signals, we find that differences in eQTL effect sizes between datasets are dominated by biological differences between cell types and tissues rather than technical differences in sample processing. Uniformly processed summary statistics provided us with a unique opportunity to characterise eQTL diversity across 69 distinct cell types and tissues. Consistent with previous analyses by the GTEx project, we find high levels of *cis*-eQTL sharing between most bulk tissues. In contrast, we find that a much smaller proportion of eQTLs are shared between purified cell types and bulk tissues, and between different cell types. This eQTL diversity also manifests itself at the level of disease colocalisation, where we detect many novel colocalisations that are missed when analysing GTEx data alone. Finally, in addition to gene expression QTLs, we have identified QTLs at the levels of exon expression, transcript usage, and splicing, which were often absent from the original studies. Our uniformly processed QTL summary statistics and fine mapping results are available from the eQTL Catalogue FTP server and REST API and they can also be explored using the Ensembl Genome Browser *(28)* (Figure 1B).

**Figure 1.**
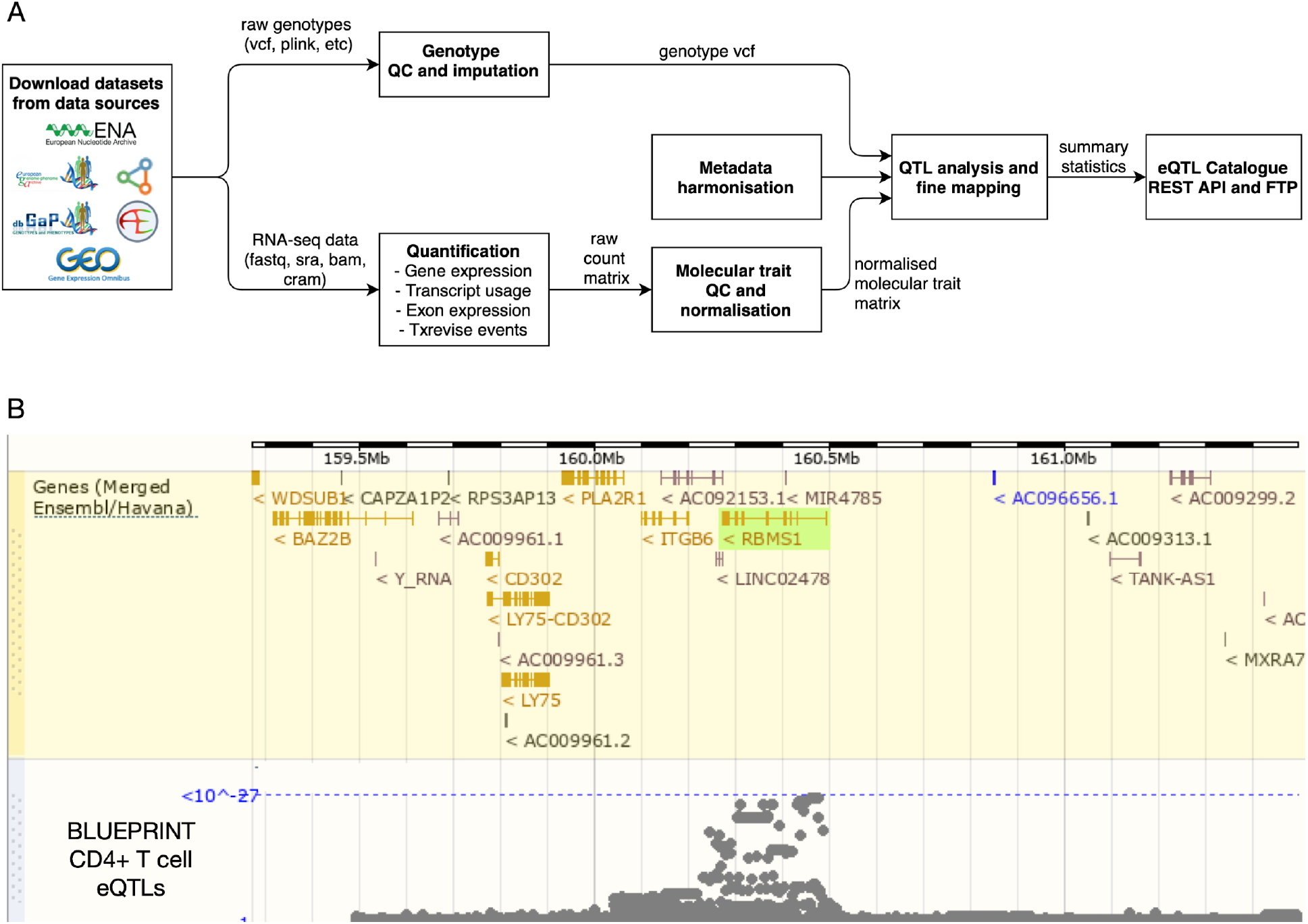
Overview of the eQTL Catalogue database. (**A**) A high-level representation of the uniform data harmonisation and eQTL mapping process. Supplementary Figure 1 provides a schematic illustration of the different quantification methods. (**B**) eQTL Catalogue summary results for the *RBMS1* gene in BLUEPRINT CD4+ T cells, viewed via the Ensembl Genome Browser.

**Figure 2.**
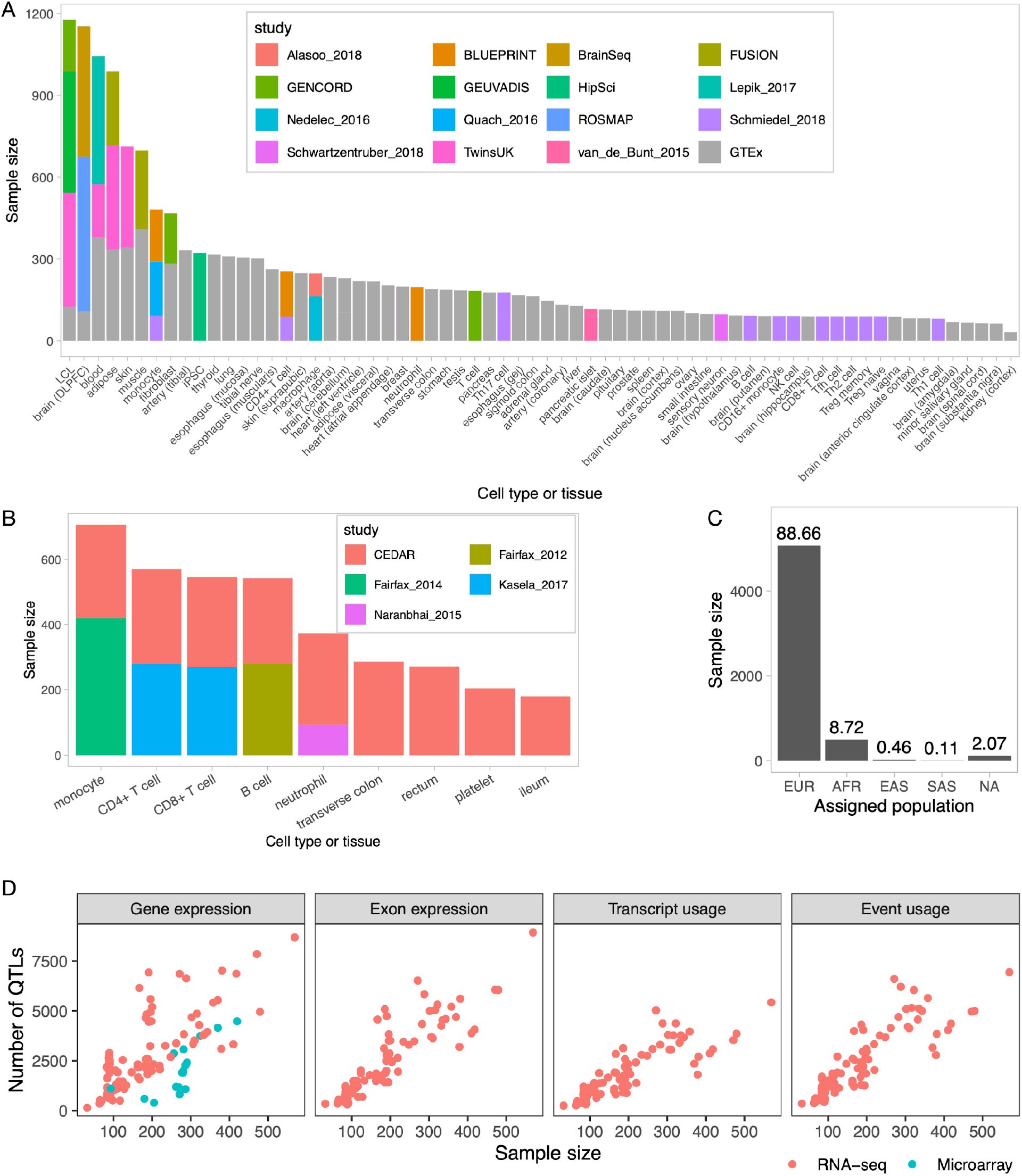
Overview of studies and samples included in the eQTL Catalogue. (**A**) Cumulative RNA-seq sample size for each cell type and tissue across 16 studies. Datasets from stimulated conditions have been excluded to improve readability. DLPFC - dorsolateral prefrontal cortex, iPSC - induced pluripotent stem cell, LCL - lymphoblastoid cell line. (**B**) The cumulative microarray sample size for each cell type and tissue across five studies. Datasets from stimulated conditions have been excluded to improve readability. (**C**) The number of unique donors assigned to the four major superpopulations in the 1000 Genomes Phase 3 reference dataset. Detailed assignment of donors to the four superpopulations in each study is presented in Supplementary Table 1. Superpopulation codes: EUR - European, AFR - African, EAS - East Asian, SAS - South Asian, NA - unassigned. (**D**) The relationship between the sample size of each dataset and the number of associations detected with each quantification method. The number of QTLs on the y-axis is defined as the number of genes with at least one significant QTL (FDR < 0.05).

## Results

### Studies, datasets and samples included in the eQTL Catalogue

We downloaded raw gene expression and genotype data from 16 RNA-seq and five microarray studies from various repositories. The RNA-seq data consisted of 17,210 samples spanning 95 datasets (defined as distinct cell types, tissues or contexts in which eQTL analysis was performed separately). These 95 datasets originated from 66 distinct cell types and tissues and ten stimulated conditions (Figure 2A). Similarly, the 4,631 microarray samples spanned 17 datasets from eight distinct cell types and tissues and three stimulated conditions (Figure 2B). While most cell types and tissues were profiled only by two of the largest studies (GTEx *(1)* and Schmiedel_2018 *(21)*, Figure 2A), 13 cell types or tissues were captured by multiple studies, allowing us to characterise both technical and biological variability between datasets and studies. The total number of unique donors across studies was 5,714, of which 89% had predominantly European ancestries and only 9% had African or African American ancestries, with other ancestries being rare (Figure 2C, Supplementary Table 1). Thus, similarly to most GWAS studies, published eQTL studies also suffer from a lack of genetic diversity *(29)*.

To uniformly process a large number of eQTL studies, we designed a modular and robust data analysis workflow (Figure 1A). First, we performed extensive quality control and imputed missing genotypes using the 1000 Genomes Phase 3 reference panel *(30)* (Supplementary Table 3). For RNA-seq datasets, we performed QTL mapping for the four molecular traits described above (Figure 1A, Supplementary Figure 1). The QTL analysis was performed separately in each dataset (i.e. separately for each cell type or tissue within each study). We found the largest number of QTLs at the level of gene expression, but for all molecular traits the number of significant associations scaled approximately linearly with the sample size (Figure 2D, Supplementary Material 1). For microarray datasets, we performed the analysis only at the gene level but found the same linear trend (Figure 2D, Supplementary Material 1). Our remaining analyses focus on the RNA-seq-based eQTL datasets as they cover a more comprehensive range of cell types and tissues, and account for most of the samples in the eQTL Catalogue.

### Biological and technical variability between studies and datasets

First, we assessed if the gene expression and eQTL signals were dominated by technical differences between studies (Supplementary Tables 2-3) rather than true biological differences between cell types and tissues. We visualised median transcripts per million (TPM) gene expression estimates from each dataset using multidimensional scaling (MDS). Reassuringly, we found that the datasets clustered predominantly by cell type or tissue of origin, rather than by studies or other technical factors (Figure 3A). Notably, except for brain tissues, whole blood and testis, most other bulk tissues had relatively similar gene expression profiles (Figure 3A). In contrast, datasets from purified cell types such as lymphoblastoid cell lines (LCLs), monocytes, neutrophils, induced pluripotent stem cells (iPSCs), and B and T lymphocytes had more distinct gene expression profiles (Figure 3A).

**Figure 3.**
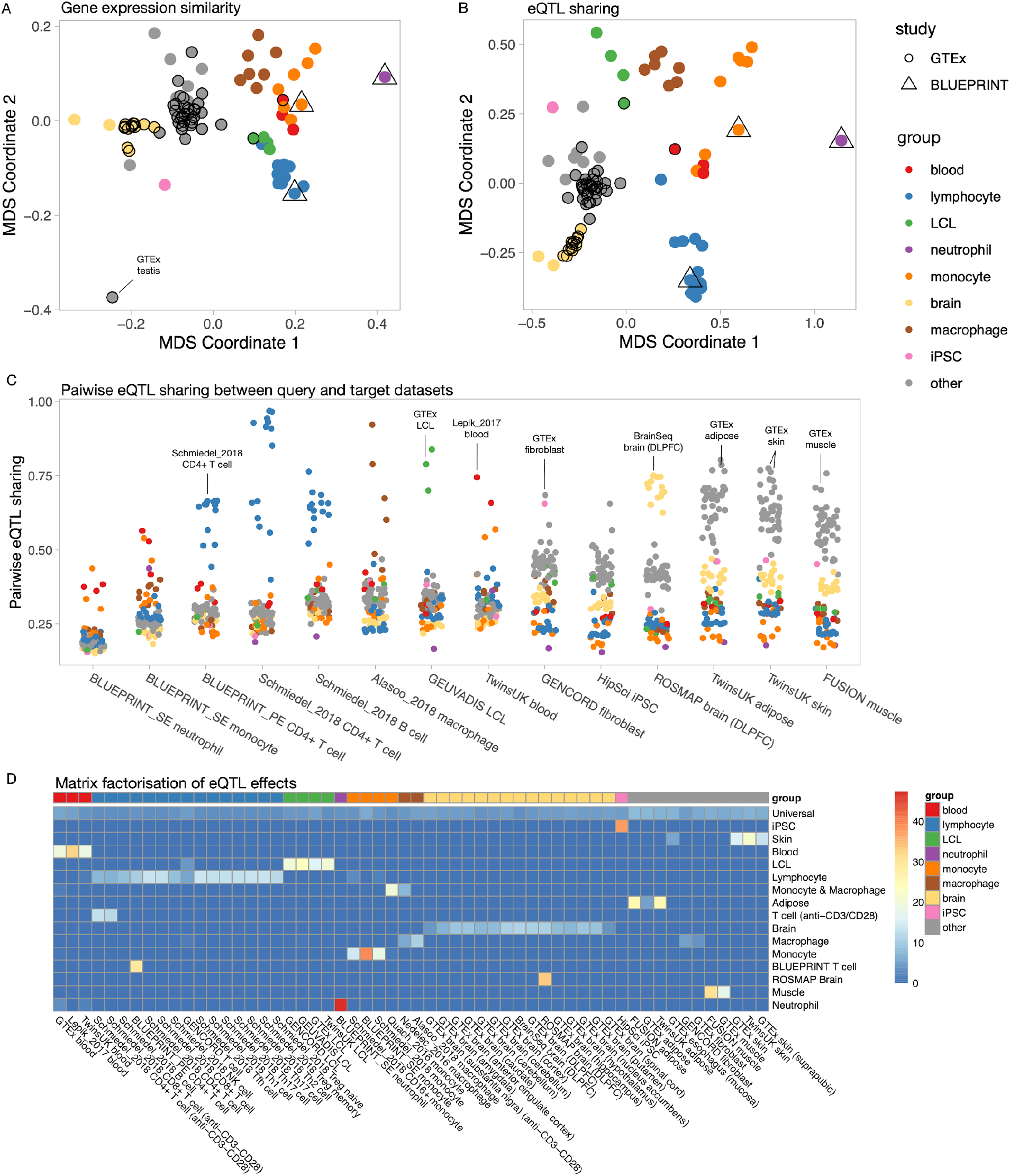
Gene expression similarity between datasets predicts eQTL similarity. (**A**) Multidimensional scaling (MDS) analysis of median gene expression across datasets. The pairwise similarity between datasets was calculated using Pearson’s correlation. Datasets from GTEx and BLUEPRINT studies have been highlighted to demonstrate that they cluster with other matching cell types and tissues. (**B**) MDS analysis of eQTL sharing across datasets. Pairwise eQTL sharing between datasets was estimated using the Mash model *(19)*. The complete matrix is presented in Supplementary Figure 2. (**C**) Visualisation of eQTL sharing estimates between selected representative tissues (x-axis) and all other cell types and tissues in the eQTL Catalogue. The individual points have been coloured according to the major cell type and tissue groups from panel A. (**D**) Matrix factorisation of the eQTL effect sizes across all eQTL Catalogue datasets. Only datasets with non-zero loading on one or more cell-type- and tissue-specific factors (excluding the universal factor) are shown.

Next, we performed the same similarity analysis on eQTL effect sizes. To overcome the high uncertainty associated with effect size estimates, especially in datasets with small sample sizes, we used the recently developed multiple adaptive shrinkage (mash) model *(19)*. Mash improves eQTL effect size estimates by sharing information both across datasets as well as individual eQTLs. We limited our analysis to 54,733 fine mapped eQTLs (see Methods) and defined two eQTLs to be shared between a pair datasets if they had the same sign and their effect sizes did not differ more than two-fold. We calculated pairwise eQTL sharing estimates for all 95 RNA-seq datasets (including 48 tissues from GTEx v7) and projected those onto two dimensions using MDS. Reassuringly, we found that if the same cell type or tissue was profiled in multiple studies, then their eQTL effect sizes often showed a high degree of concordance (Figure 3B, Supplementary Figures 2-3). For example, LCLs from TwinsUK, GENCORD and GEUVADIS clustered together with LCLs from GTEx (Figure 3B) and exhibited median sharing of ~80% (Figure 3C, Supplementary Figures 2-3). The same was also true for the brain (GTEx, ROSMAP and BrainSeq studies), whole blood (GTEx, TwinsUK and Lepik_2017 studies), muscle (GTEx and FUSION), skin (GTEx and TwinsUK) and adipose tissues (GTEx, TwinsUK and FUSION), which all had median intra-tissue sharing of ~70% (Figure 3C). Moreover, the two-dimensional MDS plot of pairwise eQTL similarity (Figure 3B) was broadly similar to the pairwise gene expression similarity plot presented above (Figure 3A), suggesting that high gene expression similarity and a high degree of eQTL sharing both reflect similarity in the underlying regulatory state of cells.

Finally, we focussed on the patterns of sharing between different cell types and tissues. We found that 46-80% (median 62%) of the eQTLs were shared between most pairs of bulk tissues (Figure 3C). The exception to this pattern were the brain tissues and whole blood that formed separate clusters in the MDS analysis (Figure 3B) and shared a median of 45% and 35% of the eQTLs with other tissues, respectively (Figure 3C). In contrast, purified immune cell types (LCLs, neutrophils, monocytes, macrophages and lymphocytes) formed distinct clusters on the MDS plot (Figure 3B) and had much lower eQTL sharing both with whole blood as well as other bulk tissues (Figure 3C). Thus, although our results reconfirm the generally high level of *cis*-eQTL sharing between bulk tissues, they also reveal a much greater *cis*-eQTL diversity between purified cell types and especially immune cells. Importantly, this diversity is missed when analysing highly tissue-focused eQTL studies such as GTEx.

### Matrix factorisation identifies cell-type- and tissue-specific latent factors shared across datasets

To better understand the eQTL sharing patterns between cell types and tissues, we turned to a recently developed semi-nonnegative sparse matrix factorisation (sn-spMF) model that can directly identify latent factors from eQTL summary statistics *(31)*. When applied to the fine mapped eQTL Catalogue summary statistics, sn-spMF detected 16 independent factors (Figure 3D). The largest universal factor was broadly shared between all datasets and accounted for ~37.5% of the independent fine mapped eQTLs (Supplementary Figure 4). The remaining 15 factors captured cell-type- and tissue-specific effects (Figure 3D). Overall, matrix factorisation identified many of the same patterns detected in the pairwise eQTL sharing analysis (Figure 2B). For example, lymphocytes, LCLs, iPSC, monocytes, macrophages, neutrophils, stimulated T cells as well as brain and blood tissues all had their individual factors. Notably, these celltype- and tissue-specific factors were shared across multiple studies (Figure 3D).

Although most eQTLs were highly shared between bulk tissues (Figure 3B-C), our factor analysis still detected independent factors capturing eQTLs that were specific to muscle, skin and adipose tissues from the FUSION *(32)*, GTEx *(1)* and TwinsUK *(33)* studies. Brain, blood, adipose, muscle and skin tissues had larger sample sizes than other bulk tissues and purified cell types (Figure 2A), allowing us to obtain more accurate eQTL effect size estimates. Thus, we expect to detect additional tissue-specific factors as the sample sizes of the respective tissues increase *(31)*. Finally, only two of the 16 factors were specific to a single dataset (BLUEPRINT CD4+ T cells and ROSMAP brain samples), suggesting that although batch effects between datasets exist, they are not a major factor confounding our analysis.

A major advantage of the matrix factorisation is that it allows us to focus on a small number of biologically meaningful factors shared between one or more datasets rather than comparing the eQTL effect sizes in 95 individual datasets. This level of summarisation is going to be increasingly important as the number of datasets included in the eQTL Catalogue increases. For example, a *cis*-eQTL for *RBMS1* had large effects in both BLUEPRINT and Schmiedel_2018 CD4+ T cell datasets and smaller significant effects in multiple other T cell subsets from Schmiedel *et al.* (Figure 4A). Consequently, the two factors with the largest loadings for this eQTL were the BLUEPRINT CD4+ T cell factor and the general lymphocyte factor (Figure 4B). The *RBMS1* eQTL also colocalised with a GWAS signal for lymphocyte count *(34)* in BLUEPRINT CD4+ T cells (PP4 = 0.94) (Figure 4C), illustrating how a lymphocyte-specific eQTL might contribute to the regulation of lymphocyte count in whole blood. Notably, we did not detect this colocalisation in any of the 49 GTEx tissues.

**Figure 4.**
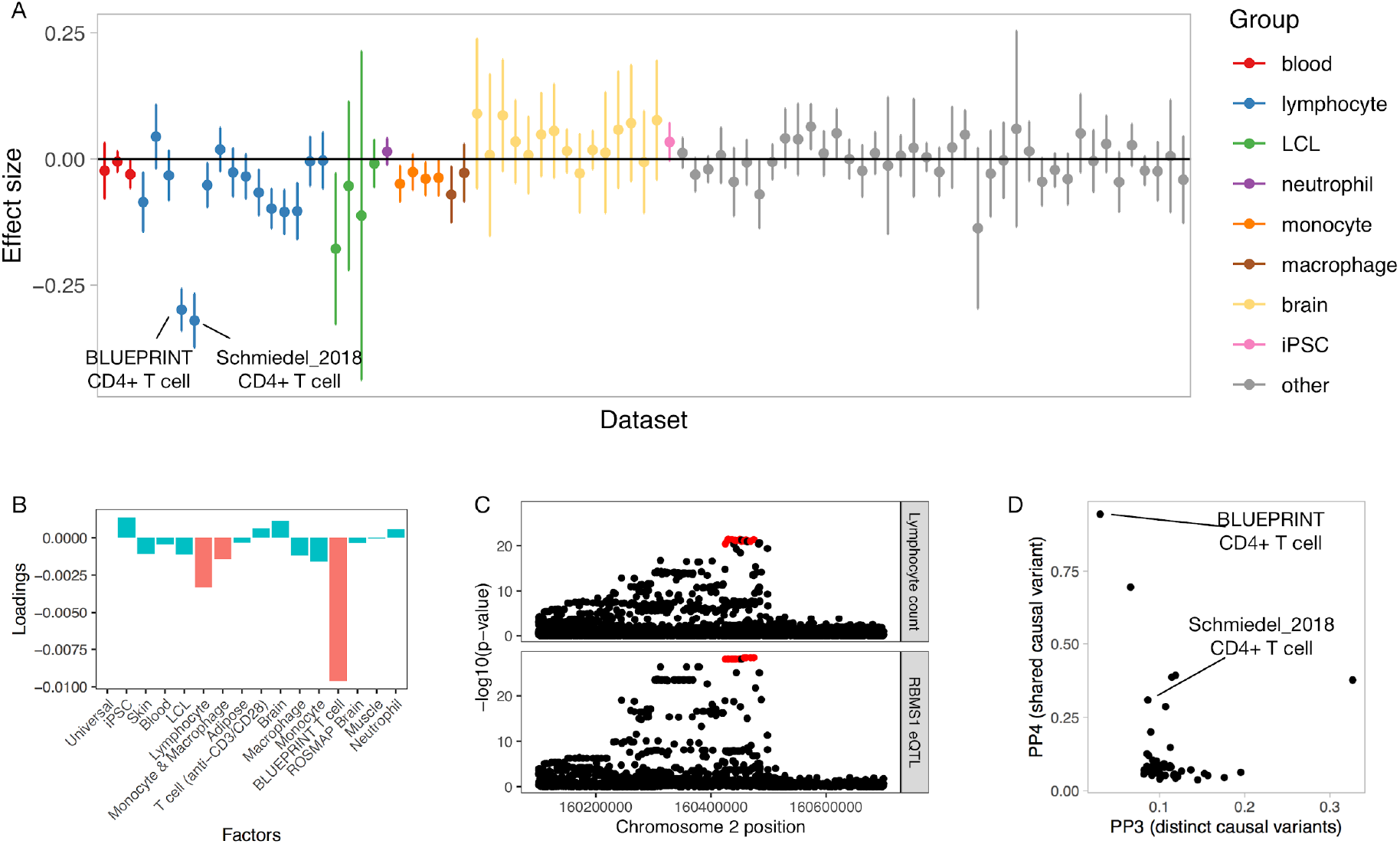
CD4+ T cell-specific eQTL at the *RBMS1* locus colocalises with a GWAS hit for lymphocyte count. (**A**) Effect sizes and 95% confidence intervals for the *RBMS1* eQTL across all eQTL Catalogue datasets (naive conditions only). (**B**) Factor loadings for the *RBMS1* lead variant (rs6753933) from the sn-spMF model. **(C**) Regional association plot for lymphocyte count (top panel) and *RBMS1* eQTL in the BLUEPRINT CD4+ T cells. The fine mapped eQTL credible set is highlighted in red. (**D**) Colocalisation posterior probabilities of a shared causal variant (PP4) and two distinct causal variants (PP3) for the fine mapped *RBMS1* lead variant (rs6753933) across all eQTL Catalogue datasets.

### eQTL Catalogue finds novel colocalisations missed in GTEx

Our eQTL sharing analysis demonstrated that the eQTL Catalogue contains many additional eQTLs not present in GTEx. To quantify how these novel eQTLs might improve the interpretation of complex trait and disease associations, we performed colocalisation between GWAS summary statistics for 14 traits and either the eQTL Catalogue datasets or all GTEx v8 tissues. To ensure that each independent GWAS locus was counted only once, we first partitioned GWAS summary statistics into approximately independent LD blocks *(35)*. Overall, we detected at least one colocalising eQTL (PP4 > 0.8) for 4,429 independent loci across 14 traits, 373 (8.4%) of which were only detected in one of the eQTL Catalogue datasets and not captured by GTEx v8 (max PP4 < 0.8). The fraction of novel colocalising loci varied from 5% for height to 14% for lupus (Supplementary Figure 5), suggesting that a substantial fraction of trait colocalisations might be missed if the analysis is only restricted to GTEx.

However, we often detected many novel colocalisations even in those eQTL Catalogue datasets that were already captured by GTEx (e.g. blood, skin, muscle, adipose and brain tissues, Figure 2A). These additional colocalisations could be either due to thresholding effects (just below or above the PP4 > 0.8 threshold), increased sample sizes in the eQTL Catalogue, and biological and population differences between datasets or other technical factors. For example, we found that the number of novel colocalisations detected for height GWAS increased linearly with the eQTL sample size with no particular dataset standing out (Figure 5A). In contrast, for some trait and eQTL dataset pairs, we detected considerably more colocalisations than we would have expected at the given sample size. For example, we observed six novel colocalisations with lymphocyte count in BLUEPRINT CD4+ T cells (including the *RBMS1* example in Figure 4C), which was three times more than in any other dataset of comparable sample size (Figure 5B).

**Figure 5.**
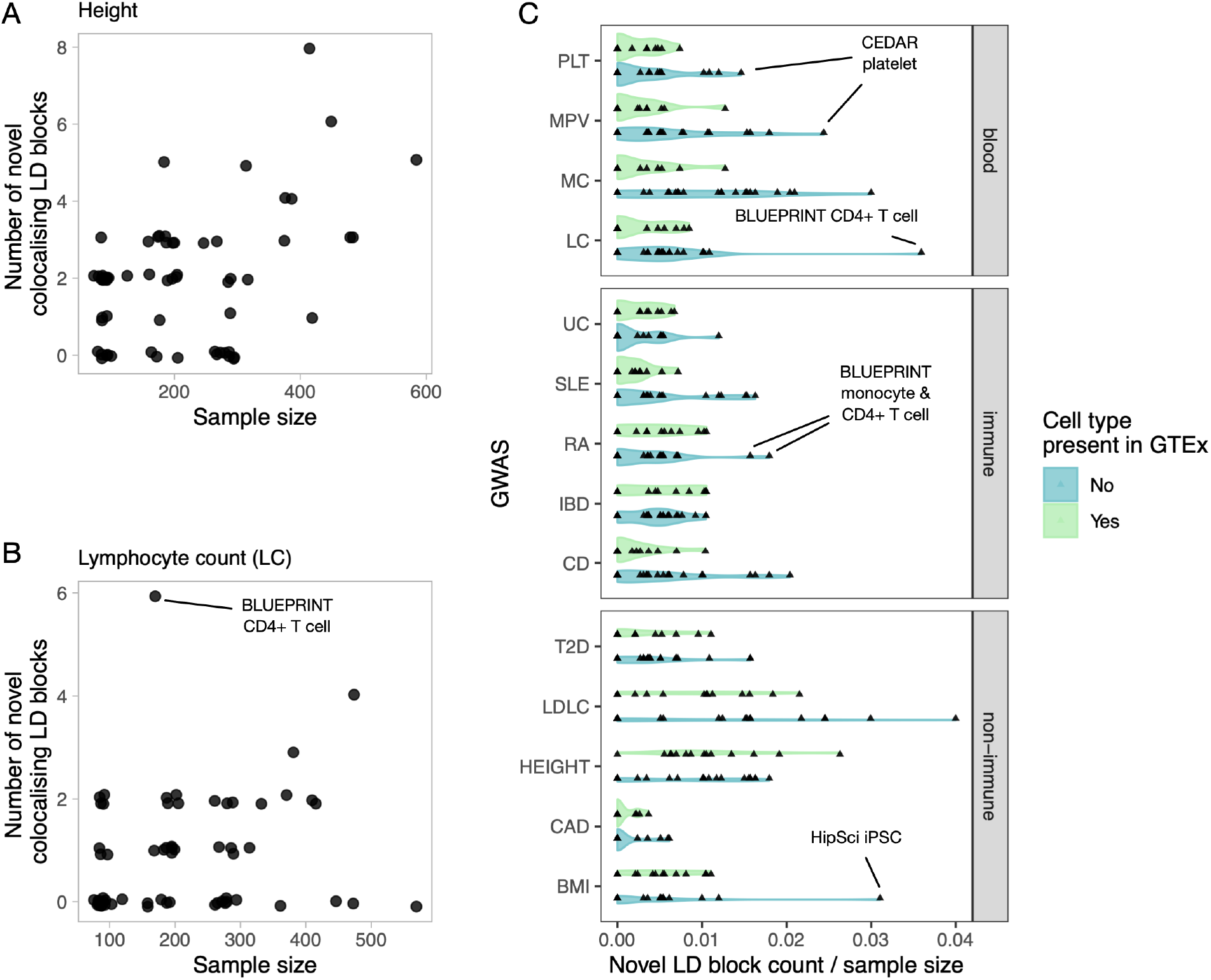
Additional GWAS colocalisations detected in the eQTL Catalogue relative to GTEx v8. (**A**) The number of novel height GWAS loci that colocalise with eQTLs in each cell type or tissues as a function of eQTL dataset size. (**B**) The number of novel lymphocyte count GWAS loci that colocalise with eQTLs in each cell type or tissues as a function of eQTL dataset size. (**C**) The number of novel colocalising loci detected for the 14 GWAS traits in each cell type and tissue from eQTL Catalogue divided by the eQTL sample sizes. The eQTL Catalogue cell types and tissues were grouped according to whether they were present in GTEx (blood, LCL, adipose, muscle, skin, brain) or not (T cells, B cells, monocytes, macrophages, neutrophils and iPSCs). GWAS traits: PLT - platelet count, MPV - mean platelet volume, MC - monocyte count, LC - lymphocyte count, UC - ulcerative colitis, SLE - systemic lupus erythematosus, RA - rheumatoid arthritis, IBD - inflammatory bowel disease, CD - Crohn’s disease, T2D - type 2 diabetes, height, CAD - coronary artery disease, BMI - body mass index, LDLC - LDL cholesterol.

To assess if some eQTL datasets were particularly relevant for specific GWAS traits, we assigned each dataset a ‘novelty score’ by dividing the number of novel colocalisations detected in that dataset by its sample size. For each GWAS trait, we then asked if the novelty scores were higher for datasets from cell types and tissues missing in GTEx compared to the datasets that were already well captured by GTEx. While there was considerable overlap between the two distributions (Figure 5C), we detected several trait-dataset pairs where the number of novel colocalisations observed was higher than expected for a given sample size. For example, the largest number of novel colocalisations for platelet count (PLT) and mean platelet volume (MPV) was detected in the CEDAR *(36)* platelet dataset (Figure 5C). Similarly, we observed most novel colocalisations for monocyte and lymphocyte count in BLUEPRINT monocyte and CD4+ T cell datasets, respectively. These results suggest that many novel colocalisations detected in the eQTL Catalogue relative to GTEx cannot be explained by sampling or technical variation alone and are likely to reflect cell-type-specific genetic effects.

### A subset of colocalisations manifest at the transcript level

Multiple studies have demonstrated that some colocalisations between QTLs and complex traits only manifest at the level of RNA splicing and transcript usage *(37, 38)*. To quantify this in the eQTL Catalogue, we performed colocalisation analysis between the 14 complex traits mentioned above and all QTLs detected with the three transcript-level quantification methods (Supplementary Figure 1). We found that 586/3394 (17.2%) colocalisations in independent LD blocks were only detected using one of the three transcript-level traits and not by traditional eQTLs in any of the 95 RNA-seq datasets (Figure 6A). However, this is likely to be underestimated because transcript and gene-level QTLs could be colocalising with independent GWAS signals within the same LD block *(1)*. Furthermore, our gene expression quantification was based on the total read count, which can also capture larger splicing changes, especially as the number of datasets and sample sizes increase.

**Figure 6.**
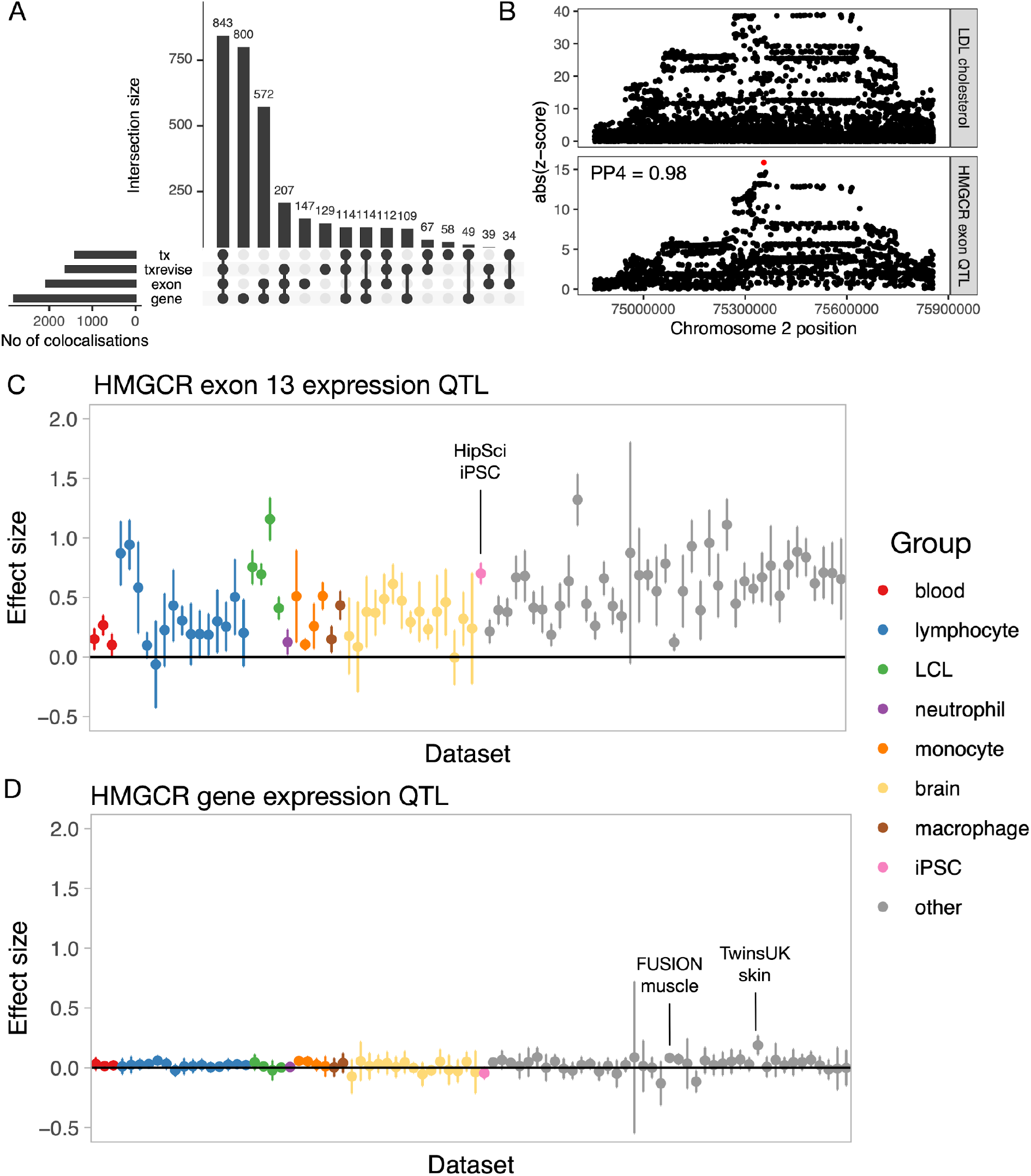
Colocalisation between transcript-level QTLs and complex traits. (**A**) Complex trait colocalisations (independent LD blocks) stratified by the quantification methods that they were detected with. In addition to gene-level eQTLs, we also used three transcript-level quantification methods (exon expression (exon), transcript usage (tx), and promoter, splicing and 3’ end usage events (txrevise)). (**B**) Regional association plot for LDL cholesterol (top panel) and *HMGCR* exon 13 QTL in the HipSci iPSC dataset. SuSiE fine mapped the exon QTL to a single intronic variant (rs3846662, represented by the red dot) which was missing from the GWAS summary statistics. (**C**) Exon 13 expression QTL effect sizes and 95% confidence intervals for the fine mapped causal variant (rs3846662) across eQTL Catalogue datasets. (**D**) Gene expression QTL effect sizes and 95%confidence intervals for the fine mapped causal variant (rs3846662) across eQTL Catalogue datasets.

To illustrate this, we looked at the colocalisation between LDL cholesterol and an exon expression QTL for *HMGCR.* The gene product of *HMGCR* is a known target for statins, and the link between exon 13 inclusion and circulating LDL cholesterol levels has been reported previously *(38, 39)*. Our analysis detected colocalisation (PP4 > 0.8) between the expression of exon 13 of the *HMGCR* gene and LDL cholesterol in 51/95 datasets. We saw the strongest association in the HipSci *(6)* induced pluripotent stem cell dataset, where we were able to fine map the exon QTL to a single causal variant (rs3846662, posterior probability = 1) (Figure 6B). The same colocalisation was also detected by transcript usage in 18/95 datasets and by txrevise in 29/95 datasets. Although the colocalisation was also seen at the level of gene expression in the FUSION *(32)* muscle dataset (PP4 = 0.99, Supplementary Figure 6), the 95% credible set contained a total of 46 variants. Furthermore, the standardised effect size of the fine mapped variant on exon expression (Figure 6C) was considerably larger than on gene expression (Figure 6D) in all datasets (Figure 6C-D). Thus, even though some transcript-level QTLs can manifest as standard eQTLs in large datasets, having access to summary statistics from different quantification methods can inform on the identity and functional impact of the causal variant as well as provide stronger genetic instruments for future Mendelian randomisation applications.

## Discussion

We believe that the main value of the eQTL Catalogue lies in the uniformly processed genelevel and transcript-level QTL summary statistics and statistical fine mapping results. We have thus sought to make the data as easy to use as possible. By mapping cell and tissue types to standard ontology terms, we make it easy to discover which studies contain the tissues and cell types of interest to the users. We have further re-imputed genotypes using the 1000 Genomes Phase 3 reference panel for all studies using genotyping microarrays, ensuring that the same set of genetic variants is present in most studies. We have used a consistent set of molecular trait identifiers (genes, exons, transcripts, events) across all datasets, ensuring that genetic effects can directly be compared across datasets (e.g. Figures 4A and 6C-D). Finally, we have released credible sets from statistical fine mapping analysis, which can help to further characterise loci with multiple independent signals and paves the way for fine-mapping-based colocalisation approaches *(25)*. We will progressively expand the resource to all accessible human datasets.

The relationship between gene expression similarity and eQTL sharing has been noticed before. For example, two studies conducted in stimulated monocytes and macrophages found that the number of differentially expressed genes between cell states correlates with the number of state-specific eQTLs *(38, 40)*. This correlation raises an exciting prospect that once a *sufficient* sample size has been reached in a given cell type or tissue, the discovery of novel eQTL can be maximised by focussing on cell types and cell states with low gene expression similarity to existing eQTL datasets. Of course, the definition of what is sufficient depends on the downstream use case of interest. While many cell-type- and tissue-specific *cis*-eQTLs can be detected with a sample size of a few hundred individuals (Figure 3D), other applications such as expression-mediated heritability analysis *(4)*, Mendelian randomisation *(18)* and *trans-eQTL* analysis *(2)* benefit from much larger sample sizes.

A limitation of our automated RNA-seq processing and eQTL mapping workflow is that we have not tailored our analyses to specific studies. For example, although the TwinsUK *(33)* and HipSci *(6)* studies collected samples from multiple related individuals, we used only a subset of samples (TwinsUK: 1,364 of 2,505 total, HipSci: 322 of 513 total) from unrelated individuals to avoid pseudoreplication when using linear regression. Similarly, for the six studies containing individuals from non-European and admixed populations (Supplementary Table 1), we jointly analysed all samples with six genotype principal components as covariates. However, stratified analyses *(41)* or approaches taking into account local ancestry *(42, 43)* might be more appropriate in this specific setting. Access to individual-level data will enable us to revisit these decisions as new analytical approaches and computational workflows become available.

To ensure that the eQTL Catalogue is a comprehensive resource that encompasses tissue and human population diversity, we encourage researchers to contribute their eQTL datasets (contact). Unfortunately, we have been unable to include some existing datasets due to consent limitations or restrictions on sharing individual-level genetic data. These limitations could be overcome in the future by federated data analysis approaches, where the eQTL analysis is performed at remote sites using our analysis workflows, and only summary statistics are shared with the eQTL Catalogue. To this end, we will continue to improve the usability and portability of our data analysis workflows and will make them available via community efforts such as the nf-core *(44)* repository.

## Methods

### Data access and informed consent

Gene expression and genotype data from two studies (GEUVADIS and CEDAR) were available for download without restrictions from ArrayExpress *(45)*. For all other datasets, we applied for access via the relevant Data Access Committees. The database accessions and contact details of the individual Data Access Committees can be found on the eQTL Catalogue website (http://www.ebi.ac.uk/eqtl/Studies/). In our applications, we explained the project and our intent to share the association summary statistics publicly. Ethical approval for the project was obtained from the Research Ethics Committee of the University of Tartu (approval 287/T-14).

### Genotype data

#### Pre-imputation quality control

We aligned the strands of the genotyped variants to the 1000 Genomes Phase 3 reference panel using Genotype Harmonizer *(46)*. We excluded genetic variants with Hardy-Weinberg p-value < 10^−6^, missingness > 0.05 and minor allele frequency < 0.01 from further analysis. We also excluded samples with more than 5% of their genotypes missing.

#### Genotype imputation and quality control

We pre-phased and imputed the genotypes to the 1000 Genomes Phase 3 reference panel *(30)* using Eagle v2.4.1 *(47)* and Minimac4 *(48)*. After imputation, we converted the coordinates of genetic variants from the GRCh37 reference genome to the GRCh38 using CrossMap v0.4.1 *(49)*. We used bcftools v1.9.0 to exclude variants with minor allele frequency (MAF) < 0.01 and imputation quality score R2 < 0.4 from downstream analysis. The genotype imputation and quality control steps are implemented in <u>eQTL-Catalogue/genimpute</u> (v20.11.1) workflow available from GitHub (see URLs).

#### Assigning individuals to reference populations

We used PLINK *(50)* v1.9.0 with ‘--indep-pairwise 50000 200 0.05’ to perform LD pruning of the genetic variants and LDAK *(51)* to project new samples to the principal components (PCs) of the 1000 Genomes Phase 3 reference panel *(30)*. To assign each genotyped sample to one of four superpopulations, we calculated the Euclidean distance in the PC space from the genotyped individual to all individuals in the reference dataset. Distance from a sample to a reference superpopulation cluster is defined as a mean of distances from the sample to each reference sample from the superpopulation cluster. We explored distances between samples and reference superpopulation clusters using different numbers of PCs and found that using 3 PCs worked best for inferring the superpopulation of a sample. Then, we assigned each sample to a superpopulation if the distance to the closest superpopulation cluster was at least 1.7 times smaller than to the second closest one (Supplementary Figure 7). We used this relatively relaxed threshold because our aim was to get an approximate estimate of the number of individuals belonging to each superpopulation. Performing a population-specific eQTL analysis would probably require a much more stringent assignment of individuals to populations. The population assignment steps are implemented in the <u>eQTL-Catalogue/qcnorm</u> (v20.12.1) workflow available from GitHub (see URLs).

### Microarray data

#### Data normalisation

All five microarray studies currently included in the eQTL Catalogue (CEDAR *(36)*, Fairfax_2012 *(52)*, Fairfax_2014 *(53)*, Kasela_2017 *(54)*, Naranbhai_2015 *(55)*) used the same Illumina HumanHT-12 v4 gene expression microarray. The database accessions for the raw data can be found on the eQTL Catalogue website (http://www.ebi.ac.uk/eqtl/Studies/). Batch effects, where applicable, were adjusted for with the function removeBatchEffect from the limma v.3.40.6 R package *(56)*. The batch adjusted log2 intensity values were quantile normalized using the lumiN function from the lumi v.2.36.0 R package *(57)*. Only the intensities of 30,353 protein-coding probes were used. The raw intensity values for the five microarray datasets have been deposited to Zenodo (doi: https://doi.org/10.5281/zenodo.3565554).

#### Detecting sample mixups

We used Genotype harmonizer *(46)* v1.4.20 to convert the imputed genotypes into TRITYPER format. We used MixupMapper *(58)* v1.4.7 to detect sample swaps between gene expression and genotype data. We detected 155 sample swaps in the CEDAR dataset, most of which affected the neutrophil samples. We also detected one sample swap in the Naranbhai_2015 dataset.

### RNA-seq data

#### Studies

eQTL Catalogue contains RNA-seq data from the following 16 studies: ROSMAP *(59)*, BrainSeq *(60)*, TwinsUK *(33)*, FUSION *(32)*, BLUEPRINT *(20, 61)*, Quach_2016 *(62)*, Schmiedel_2018 *(21)*, GENCORD *(63)*, GEUVADIS *(64)*, Alasoo_2018 *(65)*, Nedelec_2016 *(66)*, Lepik_2017 *(67)*, HipSci *(6)*, van_de_Bunt_2015 *(68)*, Schwartzentruber_2018 *(69)*, GTEx v7 *(1)*.

#### Pre-processing

For each study, we downloaded the raw RNA-seq data from one of the six databases (European Genome-phenome Archive (EGA), European Nucleotide Archive (ENA), Array Express, Gene Expression Omnibus (GEO), Database of Genotypes and Phenotypes (dbGaP), Synapse). If the data were already in fastq format, then we proceeded directly to quantification. If the raw data were shared in BAM or CRAM format, we used the samtools collate command *(70)* to collate paired-end reads and then used samtools fastq command with ‘-F 2816 -c 6’ flags to convert the CRAM or BAM files to fastq. Since samples from GEO and dbGaP were stored in SRA format, we used the fastq-dump command with ‘--splitfiles --gzip --skip-technical --readids --dumpbase --clip’ flags to convert those to fastq. The pre-processing scripts are available from the <u>eQTL-Catalogue/rnaseq</u> GitHub repository (see URLs).

#### Quantification

We quantified transcription at four different levels: (1) gene expression, (2) exon expression, (3) transcript usage and (4) transcriptional event usage (Supplementary Figure 1). Quantification was performed using a custom Nextflow *(71)* workflow that we developed by adding new quantification methods to nf-core/rnaseq pipeline *(44)*. Before quantification, we used Trim Galore v0.5.0 to remove sequencing adapters from the fastq files.

For gene expression quantification, we used HISAT2 v2.1.0 *(72)* to align reads to the GRCh38 reference genome (Homo_sapiens.GRCh38.dna.primary_assembly.fa file downloaded from Ensembl). We counted the number of reads overlapping the genes in the GENCODE V30 *(73)* reference transcriptome annotations with featureCounts v1.6.4 *(74)*. To quantify exon expression, we first created an exon annotation file (GFF) using GENCODE V30 reference transcriptome annotations and dexseq_prepare_annotation.py script from the DEXSeq *(75)* package. We then used the aligned RNA-seq BAM files from the gene expression quantification and featureCounts with flags ‘-p -t exonic_part -s ${direction} -f -O’ to count the number of reads overlapping each exon.

We quantified transcript and event expression with Salmon v0.13.1 *(76)*. For transcript quantification, we used the GENCODE V30 (GRCh38.p12) reference transcript sequences (fasta) file to build the Salmon index. For transcriptional event usage, we downloaded precomputed txrevise *(38)* alternative promoter, splicing and alternative 3’ end annotations corresponding to Ensembl version 96 from Zenodo (https://doi.org/10.5281/zenodo.3232932) in GFF format. We then used gffread *(77)* to generate fasta sequences from the event annotations and built Salmon indices for each event set as we did for transcript usage. Finally, we quantified transcript and event expression using salmon quant with ‘--seqBias --useVBOpt --gcBias --libType’ flags. All expression matrices were merged using csvtk v0.17.0. All of these quantification methods are implemented in the <u>eQTL-Catalogue/rnaseq</u> workflow available from GitHub (see URLs). Our reference transcriptome annotations are available from Zenodo (https://doi.org/10.5281/zenodo.3366280).

#### Detecting outliers from gene expression data

The quality of the RNA-seq samples was assessed using the gene expression counts matrix. In all downstream analyses, we only included 35,367 protein-coding and non-coding RNA genes belonging to one of the following Ensembl gene types: lincRNA, protein_coding, IG_C_gene, IG_D_gene, IG_J_gene, IG_V_gene, TR_C_gene, TR_D_gene, TR_J_gene, TR_V_gene, 3prime_overlapping_ncrna, known_ncrna, processed_transcript, antisense, sense_intronic, sense_overlapping. For PCA and MDS analyses, we first filtered out invalid gene types (23,458) and genes on the sex chromosomes (1,247), TPM normalised *(78)* the gene counts, filtered out genes having median normalised expression value less than 1 and log2 transformed the matrix. We performed principal component analysis with the prcomp R package (center = true, scale = true). For multidimensional scaling (MDS) analysis, we used the isoMDS method from the MASS R package with k=2 dimensions. As a distance metric for isoMDS, we used 1 - Pearson’s correlation as recommended previously *(79)*. We plotted these two-dimensional scatter plots to visually identify outliers (Supplementary Figure 8A-B).

#### Sex-specific gene expression analysis

Previous studies have successfully used the expression of *XIST* and Y chromosome genes to ascertain the genetic sex of RNA samples *(80)*. In our analysis, we extracted all protein-coding genes from the Y chromosome, and the *XIST* gene (ENSG00000229807) expression values and TPM normalised them. Then, we calculated the mean expression level of the genes on the Y chromosome. Finally, we plotted the log2 of *XIST* expression level (X-axis) against the mean expression level of the genes on the Y chromosome (Y-axis) (Supplementary Figure 8C). In addition to detecting samples with incorrectly labelled genetic sex, this analysis also allowed us to identify cross-contamination between samples *(XIST* and Y chromosome genes expressed simultaneously, Supplementary Figure 8C).

#### Concordance between genotype data and RNA-seq samples

We used the Match Bam to VCF (MBV) method from QTLtools *(81)* which directly compares the sample genotypes in VCF format to an aligned RNA-seq BAM file. MBV can detect sample swaps, multiple samples from the same donor, and cross-contamination between RNA-seq samples. In some cases, such cross-contamination was confirmed by both the sex-specific gene expression and MBV analyses (Supplementary Figure 8D).

#### Normalisation

We filtered out samples which failed the QC step. We normalised the gene and exon-level read counts using the conditional quantile normalisation (cqn) R package v1.30.0 *(82)* with gene or exon GC nucleotide content as a covariate. We downloaded the gene GC content estimates from Ensembl biomaRt and calculated the exon-level GC content using bedtools v2.19.0 *(83)*. We also excluded lowly expressed genes, where 95 per cent of the samples within a dataset had TPM-normalised expression less than 1. To calculate transcript and transcriptional event usage values, we obtained the TPM normalised transcript (event) expression estimates from Salmon. We then divided those transcript (event) expression estimates by the total expression of all transcripts (events) from the same gene (event group). Subsequently, we used the inverse normal transformation to standardise the transcript and event usage estimates. Normalisation scripts together with containerised software are publicly available at https://github.com/eQTL-Catalogue/qcnorm.

### Metadata harmonisation

We mapped all RNA-seq and microarray samples to a minimal metadata model. This included consistent sample identifiers, information about the cell type or tissue of origin, biological context (e.g. stimulation), genetic sex, experiment type (RNA-seq or microarray) and properties of the RNA-seq protocol (paired-end vs single-end; stranded vs unstranded; poly(A) selection vs total RNA). To ensure that cell type and tissue names were consistent between studies and to facilitate easier integration of additional studies, we used Zooma (https://www.ebi.ac.uk/spot/zooma/) to map cell and tissue types to a controlled vocabulary of ontology terms from Uber-anatomy ontology (Uberon) *(84)*, Cell Ontology *(85)* or Experimental Factor Ontology (EFO) *(86)*. We opted to use an *ad-hoc* controlled vocabulary to represent biological contexts as those often included terms and combinations of terms that were missing from ontologies.

### Association testing

We performed association testing separately in each dataset and used a +/- 1 megabase *cis* window centred around the start of each gene. First, we excluded molecular traits with less than five genetic variants in their *cis* window, as these were likely to reside in regions with low genotyping coverage. We also excluded molecular traits with zero variance across all samples and calculated phenotype principal components using the prcomp R stats package (center = true, scale = true). We calculated genotype principal components using plink2 v1.90b3.35. We used the first six genotype and phenotype principal components as covariates in QTL mapping. We calculated nominal eQTL summary statistics using the GTEx v6p version of the FastQTL *(87)* software (https://github.com/francois-a/fastqtl) that also estimates standard errors of the effect sizes. We used the ‘--window 1000000 --nominal 1’ flags to find all associations in 1 Mb *cis* window. For permutation analysis, we used QTLtools v1.1 *(88)* with ‘--window 1000000 --permute 1000 --grp-best’ flags to calculate empirical p-values based on 1000 permutations. The ‘--grp-best’ option ensured that the permutations were performed across all molecular traits within the same ‘group’ (e.g. multiple probes per gene in microarray data or multiple transcripts or exons per gene in the exon-level and transcript-level analysis) and the empirical p-value was calculated at the group level. The steps described above are implemented in the <u>eQTL-Catalogue/qtlmapv</u> 20.07.2 Nextflow workflow available from GitHub (see URLs).

### Statistical fine mapping

We performed QTL fine mapping using the Sum of Single Effects Model (SuSiE) *(23)* implemented in the susieR v0.9.0 R package. We converted the genotypes from VCF format to a tabix-indexed dosage matrix with bcftools v1.10.2. We imported the genotype dosage matrix into R using the Rsamtools v1.34.0 R package. We used the same normalised molecular trait matrix used for QTL mapping and further applied a rank-based inverse normal transformation to each molecular trait to ensure that they were normally distributed. We regressed out the first six phenotype and genotype PCs separately from the phenotype and genotype matrices. We performed fine mapping with the following parameters: L = 10, estimate_residual_variance = TRUE, estimate_prior_variance = TRUE, scaled_prior_variance = 0.1, compute_univariate_zscore = TRUE, min_abs_corr = 0. Finally, we extracted the 95% credible sets and the 95% posterior inclusion probabilities for each variant belonging to the credible set. The steps described above are implemented in the <u>eQTL-Catalogue/susie-workflow</u> v20.08.3 Nextflow workflow available from GitHub (see URLs).

### Quantifying eQTL sharing between tissues, cell types and conditions

#### Identifying independent signals based on fine mapping

We extracted independent signals from the variants included in fine-mapped credible sets. At first, we selected credible sets with less than 50 variants in size and with a univariate z-score of at least 3. For every gene, we built connected components of credible sets to represent independent signals. From every connected component, we picked the lead variant – the variant with the smallest p-value across all eQTL datasets. As a result, 54,733 eQTLs remained.

#### Calculating Spearman’s correlation

We aggregated the eQTL data into a matrix of effect sizes, where each row represents a lead variant and each column an eQTL dataset. We noticed that this matrix contained many missing values. While most of the missing values were caused by the gene not being expressed in a particular cell type or tissue, some of the missing values were also caused by low allele frequency or low imputation quality score. Thus, we substituted all missing values with 0. We then calculated pairwise Spearman’s correlation between the columns of the matrix to estimate the eQTL similarity between datasets.

#### Running Mash

As an alternative to Spearman’s correlation, we used the multiple adaptive shrinkage (Mash) *(19)* model to estimate the pairwise sharing of eQTLs between datasets. Betas and standard errors of lead effects were input to the Mash model as Bhat and Shat. We set missing eQTL effect sizes to 0 and standard errors to 1. The model was fitted with alpha = 1 (exchangeable effects model). To find candidate covariance matrices, we discovered strong effects that are significant in at least one dataset with the get_significant_results method. Then we performed PCA on identified strong effects to obtain covariance matrices with cov_pca function and applied extreme deconvolution to them with cov_ed. Resulting matrices were set as a candidate covariance matrices into the model fitting. We estimated pairwise eQTL sharing between datasets with get_pairwise_sharing method by magnitude (factor of 0.5) and sign of posterior effect estimates.

### Factor analysis

We performed factor analysis using the semi-nonnegative sparse matrix factorisation (sn-spMF) model *(31)*. We included the 54,733 independent gene-variant pairs detected using statistical fine mapping (see above). The input files contained effect sizes and standard errors as reciprocal of weights of lead effects. The missing values made up 27% of the input effect size matrix. If the effect size estimate was missing in a given cell type or tissue then the effect size and weight were set to zero. To find hyper-parameter K (initial number of factors), and regularization parameters alpha and lambda, we performed a two-level grid search. In the first level, K was set to 20, 30, 40, 50, lambda and alpha were set in a range of 800 to 1800 with optimisation number of iterations = 10. In the second level, we fine-tuned the parameters by narrowing the search space to those values that lead to higher sparsity of the loading and factor matrices in the first level. At the second level, we ran the parameter optimization for 50 iterations. We picked the final matrix with a very high cophenetic coefficient (0.99) and 16 factors.

### Colocalisation

We performed colocalisation analysis on QTLs in the eQTL Catalogue against GWAS summary statistics from 14 studies downloaded from the IEU OpenGWAS database in VCF format *(89, 90)*. Our analysis included summary statistics for inflammatory bowel disease (IBD) and its two subtypes (Crohn’s disease (CD) and ulcerative colitis (UC)) *(91)*; rheumatoid arthritis (RA) *(92)*, systemic lupus erythematosus (SLE) *(93)*, type 2 diabetes (T2D) *(94)*, coronary artery disease (CAD) *(95)*, LDL cholesterol *(96)*, four blood cell type traits (lymphocyte count (LC), monocyte count (MC), platelet count (PLT), mean platelet volume (MPV)) *(34)* and two anthropometric traits (height, body mass index (BMI)) from the UK Biobank *(96)*. The variant coordinates of the GWAS summary statistics were lifted to the GRCh38 reference genome using CrossMap *(49)*. Allele frequencies of variants in five of the GWAS (IBD, CD, UC, RA, SLE) were extracted from the 1000 Genomes Phase 3 reference panel *(30)*. For all eQTL and GWAS dataset pairs, we performed colocalisation in a ± 200,000 window around each of the 54,733 fine mapped eQTL credible set lead variants (see fine mapping above). This ensured that colocalisation was also performed separately for multiple independent eQTLs of the same gene and colocalisation results were obtained in datasets in which no significant eQTL was detected for a particular gene. However, since we did not use masking or conditional analysis, many secondary eQTL colocalisations could still have been missed *(18, 97)*. Since transcript usage, exon expression and txrevise contained many more redundant phenotypes (e.g. multiple exons of the same gene), we limited colocalisation analysis for those molecular traits to the significant lead QTL variants in each dataset only (FDR < 0.01), using the same ± 200,000 *cis* window as above. We used version 3.1 of the coloc R package *(98)*. All analysis steps are implemented in the <u>eQTL-Catalogue/colocalisation</u> (v20.11.1) workflow (see URLs).

#### Quantification of novel colocalisations at the transcript level

We only included QTL and complex trait pairs with strong evidence of colocalisations (PP4 > 0.8) in our analysis. Inspired by the study by Barbeira *et al. (3)*, we summarised colocalisations at the level of approximately independent LD blocks *(35)*. Positions of approximately independent LD blocks were obtained from Berisa and Pickrell *(35)* and converted to GRCh38 coordinates using CrossMap *(49)*. If the colocalisation *cis* window overlapped two or more LD blocks, then the colocalising QTL was assigned to the LD block where the QTL lead variant was located. The number of LD blocks for which we detected at least one colocalising QTL with each quantification method was visualised using the upsetR R package *(99)*.

#### Comparative analysis with GTEx V8

Current version of the eQTL Catalogue (release 3) contains two versions of the GTEx summary statistics: uniformly processed summary statistics from GTEx v7 and the official GTEx v8 summary statistics downloaded from Google Cloud (gs://gtex-resources/GTEx_Analysis_v8_QTLs/GTEx_Analysis_v8_eQTL_all_associations). Since the sample size of GTEx v8 is approximately two times larger than GTEx v7, we decided to use the official GTEx v8 summary statistics in our comparative colocalisation analysis. This ensured that we were as conservative as possible when identifying novel colocaliations. For each GWAS trait, we summarised colocalisation signals at the level of independent LD blocks and defined an LD block to harbour a novel colocalisation signal if there was no colocalisation detected within that LD block in any of the GTEx v8 tissues. We further excluded datasets with small sample sizes (n < 150) due to their low power to detect colocalisations.

## URLs

Data analysis workflows:

- RNA-seq quantification: https://github.com/eQTL-Catalogue/rnaseq
- Normalisation and QC: https://github.com/eQTL-Catalogue/qcnorm
- Genotype imputation: https://github.com/eQTL-Catalogue/genimpute
- Association testing: https://github.com/eQTL-Catalogue/qtlmap
- Statistical fine mapping: https://github.com/eQTL-Catalogue/susie-workflow
- Colocalisation: https://github.com/eQTL-Catalogue/colocalisation

Example use cases:

- Accessing eQTL Catalogue summary statistics with tabix: https://github.com/eQTL-Catalogue/eQTL-Catalogue-resources/blob/master/tutorials/tabix_use_case.md
- Python example for querying the HDF5 files: https://github.com/eQTL-Catalogue/eQTL-SumStats/blob/master/querying hdf5 basics.ipynb

## Data availability

All eQTL Catalogue summary statistics are available under the Creative Commons Attribution 4.0 International License. The full association summary statistics and fine mapped credible sets in HDF5 and TSV format can be downloaded from the eQTL Catalogue website (https://www.ebi.ac.uk/eqtl/Data_access/). Slices of the TSV files can be accessed using tabix *(100)* and seqminer *(101)*. All of the summary statistics are also available via the REST API (https://www.ebi.ac.uk/eqtl/api-docs/). Fine mapped credible sets can be browsed using our interactive web interface (https://elixir.ut.ee/eqtl/). Database accessions for the raw gene expression and genotype datasets are listed on the eQTL Catalogue website (https://www.ebi.ac.uk/eqtl/Studies/). Our summary statistics have also been integrated into third party services such as the Open Targets Genetics Portal *(102)* and FUMA *(13)*. The gene expression matrices will be made available via the EMBL-EBI Expression Atlas *(103)*.

## Author contributions

NK and KA developed the data analysis and quality control workflows and performed quality control of the data. NK processed the RNA-seq datasets and performed the QTL analysis. JH developed and implemented the eQTL Catalogue API. JM processed the gene expression data for the Expression Atlas. LK performed microarray gene expression data normalisation and quality control. KP and MS developed the initial version of the population assignment workflow.

KP performed eQTL similarity and matrix factorisation analyses. MPS connected the Ensembl display to the eQTL Catalogue. IK created the interactive credible set browser. TB, SJ, HP, AY, ST, IP, DZ and KA supervised the work. NK and KA wrote the manuscript with input from all authors.

## Acknowledgements

The RNA-seq quantification and QTL analyses were performed at the High Performance Computing Center, University of Tartu. We thank Eleri Pihlapuu from the Grant Office of the University of Tartu, and Holly Foster and Paris Litterick from Open Targets for assistance in setting up data access agreements. We thank Jeremy Schwartzentruber, Emily Steed, Silva Kasela and Urmo Võsa for their helpful comments on the manuscript; Masahiro Kanai, Jacob Ulirsch and Hilary Finucane for feedback on the fine mapping workflow; Daniel Gaffney for guidance in setting up this project.

## Funding for datasets in the eQTL Catalogue

### BLUEPRINT

This study makes use of data generated by the Blueprint Consortium. A full list of the investigators who contributed to the generation of the data is available from www.blueprint-epigenome.eu. Funding for the project was provided by the European Union’s Seventh Framework Programme (FP7/2007-2013) under grant agreement no 282510 - BLUEPRINT.

### Fairfax_2012, Fairfax_2014 and Naranbhai_2015

Funding for the project was provided by the Wellcome Trust under awards Grants 088891 [B.P.F.], 074318 [J.C.K.] and 075491/Z/04 to the core facilities at the Wellcome Trust Centre for Human Genetics, the European Research Council under the European Union’s Seventh Framework Programme (FP7/2007-2013) (281824 to J.C.K.), the Medical Research Council (98082, J.C.K.) and the National Institute for Health Research (NIHR) Oxford Biomedical Research Centre.

### TwinsUK

TwinsUK is funded by the Wellcome Trust, Medical Research Council, European Union, the National Institute for Health Research (NIHR)-funded BioResource, Clinical Research Facility and Biomedical Research Centre based at Guy’s and St Thomas’ NHS Foundation Trust in partnership with King’s College London.

### BrainSeq

This research was supported by the Intramural Research Program of the NIMH (NCT00001260, 900142).

### Schmiedel_2018

This work was funded by the William K. Bowes Jr Foundation (P.V.) and NIH grants R24AI108564 (P.V., B.P., A.R., M.K.), S10RR027366 (BD FACSAria II), and S10OD016262 (Illumina HiSeq 2500).

### ROSMAP

Study data were provided by the Rush Alzheimer’s Disease Center, Rush University Medical Center, Chicago. Data collection was supported through funding by NIA grants P304G10161, R014G15819, R014G17917, R01AG30!46, R014G36836, U014G32984, U014G46152, the Illinois Department of Public Health, and the Translational Genomics Research Institute.

### GENCORD

Emmanouil T Dermitzakis was supported by grants from the European Research Council (260927), Swiss National Science Foundation (31003A_130342, CRSI33_130326) Louis-Jeantet Foundation, and the Blueprint Consortium. Stylianos E Antonarakis was supported by grants from the European Research Council (249968), Swiss National Science Foundation (144082), and the Blueprint Consortium.

### van_de_Bunt_2OI5

MvdB is supported by a Novo Nordisk postdoctoral fellowship run in partnership with the University of Oxford. ALG is a Wellcome Trust Senior Research Fellow in Basic Biomedical Science (095010/Z/10/Z). MIM is a Wellcome Trust Senior Investigator (WT098381) and a National Institute of Health Research Senior Investigator. PEM holds the Canada Research Chair in Islet Biology. This work was supported in part in Oxford, UK, by grants from the Medical Research Council (MRC; MR/L020149/1) and National Institutes of Health (NIH; R01 MH090941), and in Edmonton, Canada, by operating grants to PEM from the Canadian Institutes of Health Research (CIHR; MOP244739) and the ADI/Johnson & Johnson Diabetes Research Fund. Human islet isolations at the Alberta Diabetes Institute IsletCore were funded by the Alberta Diabetes Foundation and the University of Alberta. The National Institute for Health Research, Oxford Biomedical Research Centre funded islet provision at the Oxford Human Islet Isolation facility. The funders had no role in study design, data collection and analysis, decision to publish, or preparation of the manuscript.

### FUSION

Support for the FUSION Tissue Biopsy Study dataset was contributed by the NHGRI intramural projects ZIAHG000024 and Z1BHG000196, NIDDK grants DK062370, DK072193, and DK099240, NHGRI grant HG003079, American Diabetes Association Pathway to Stop Diabetes Grant 1-14-INI-07, and grants from the Academy of Finland.

## Supplementary Materials

**Supplementary Table 1.**
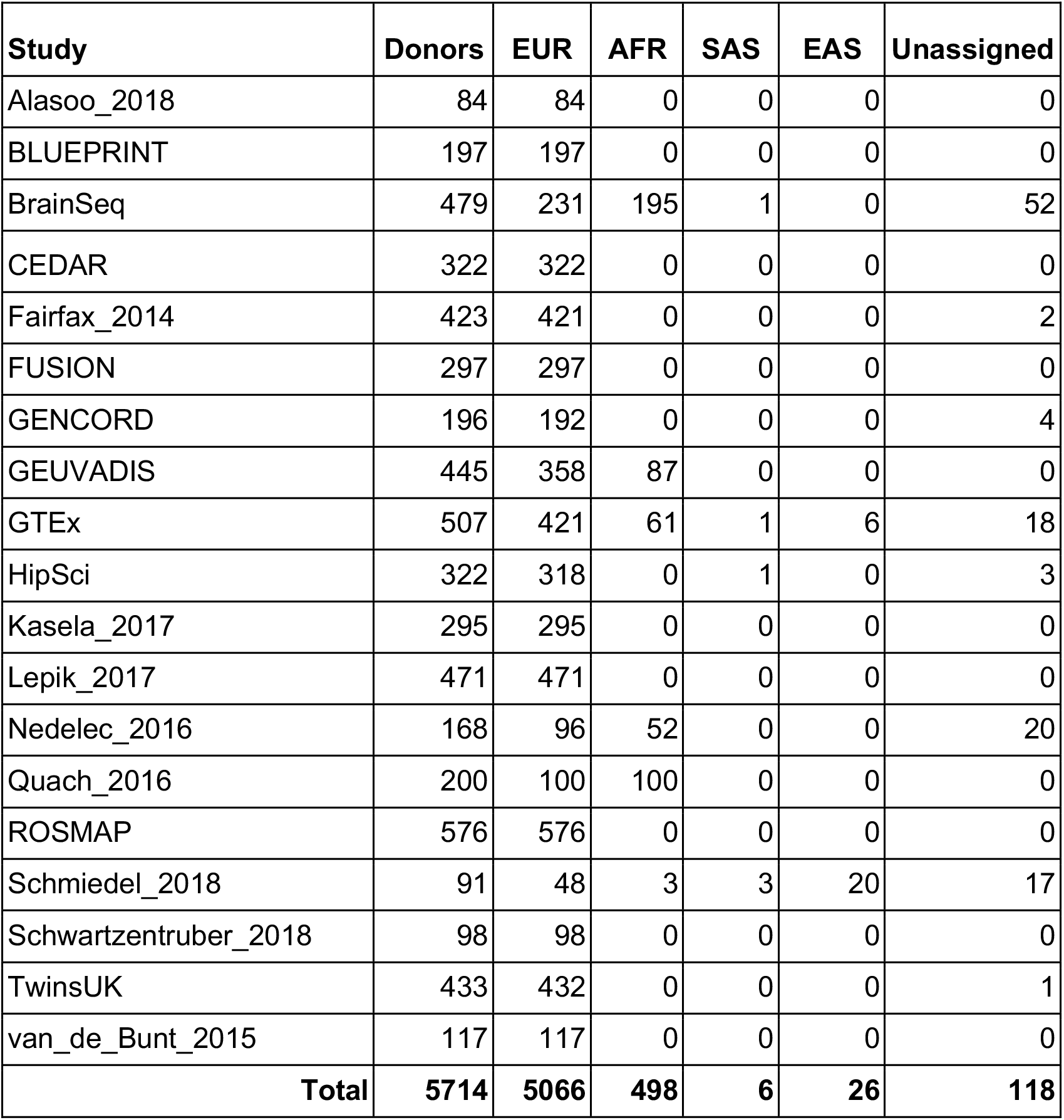
Samples assigned to the 1000 Genomes Phase 3 reference populations in each study. Note that three studies based on HipSci samples (HipSci, Alasoo_2018, Schwartzentruber_2018) and two studies based on Estonian Biobank samples (Kasela_2017, Lepik_2017) share a subset of donors by design. Furthermore, Fairfax_2012 and Naranbhai_2015 studies have been excluded because donors in these two studies are a subset of donors in Fairfax_2014. Thus, the total number of donors (n = 5,714) in this table slightly exceeds the number of unique donors. Superpopulation codes: EUR - European, AFR - African, SAS - South Asian, EAS - East Asian.

**Supplementary Table 2.**
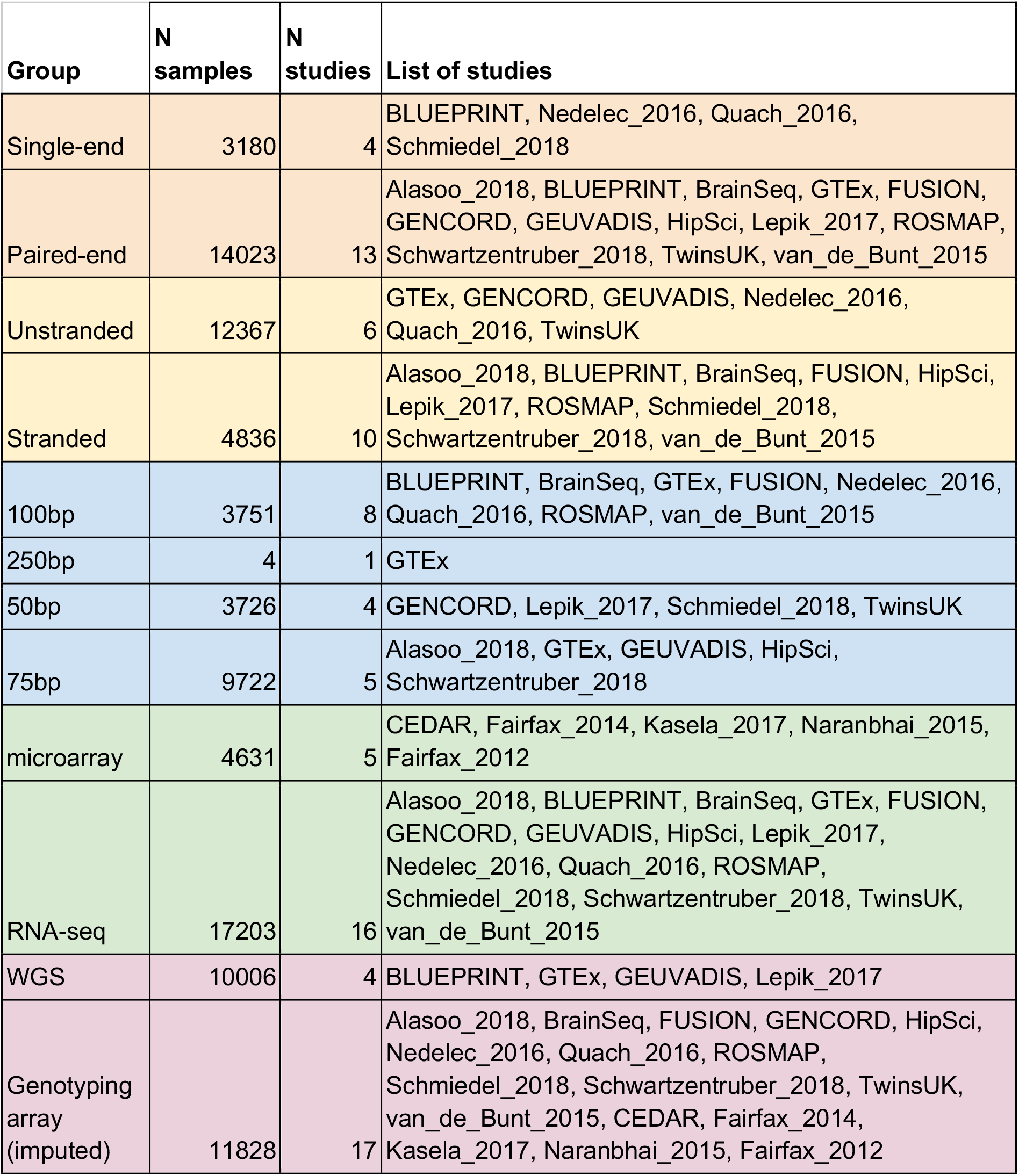
Overview of the transcriptomic samples included in the eQTL Catalogue. The samples have been classified according to RNA-seq type (single-end vs paired-end), strandedness (unstranded vs stranded), read length (50bp, 75bp, 100bp, 250bp), assay type (microarray vs RNA-seq) and genotype data type (whole-genome sequencing (WGS) vs genotyping array). Only genotyping array samples have been re-imputed by us.

| Group | N samples | N studies | List of studies |
| --- | --- | --- | --- |
| Single-end | 3180 | 4 | BLUEPRINT, Nedelec_2016, Quach_2016, Schmiedel_2018 |
| Paired-end | 14023 | 13 | Alasoo_2018, BLUEPRINT, BrainSeq, GTEx, FUSION, GENCORD, GEUVADIS, HipSci, Lepik_2017, ROSMAP, Schwartzentruber_2018, TwinsUK, van_de_Bunt_2015 |
| Unstranded | 12367 | 6 | GTEx, GENCORD, GEUVADIS, Nedelec_2016, Quach_2016, TwinsUK |
| Stranded | 4836 | 10 | Alasoo_2018, BLUEPRINT, BrainSeq, FUSION, HipSci, Lepik_2017, ROSMAP, Schmiedel_2018, Schwartzentruber_2018, van_de_Bunt_2015 |
| 100bp | 3751 | 8 | BLUEPRINT, BrainSeq, GTEx, FUSION, Nedelec_2016, Quach_2016, ROSMAP, van_de_Bunt_2015 |
| 250bp | 4 | 1 | GTEx |
| 50bp | 3726 | 4 | GENCORD, Lepik_2017, Schmiedel_2018, TwinsUK |
| 75bp | 9722 | 5 | Alasoo_2018, GTEx, GEUVADIS, HipSci, Schwartzentruber_2018 |
| microarray | 4631 | 5 | CEDAR, Fairfax_2014, Kasela_2017, Naranbhai_2015, Fairfax_2012 |
| RNA-seq | 17203 | 16 | Alasoo_2018, BLUEPRINT, BrainSeq, GTEx, FUSION, GENCORD, GEUVADIS, HipSci, Lepik_2017, Nedelec_2016, Quach_2016, ROSMAP, Schmiedel_2018, Schwartzentruber_2018, TwinsUK, van_de_Bunt_2015 |
| WGS | 10006 | 4 | BLUEPRINT, GTEx, GEUVADIS, Lepik_2017 |
| Genotyping array (imputed) | 11828 | 17 | Alasoo_2018, BrainSeq, FUSION, GENCORD, HipSci, Nedelec_2016, Quach_2016, ROSMAP, Schmiedel_2018, Schwartzentruber_2018, TwinsUK, van_de_Bunt_2015, CEDAR, Fairfax_2014, Kasela_2017, Naranbhai_2015, Fairfax_2012 |

**Supplementary Table 3.**
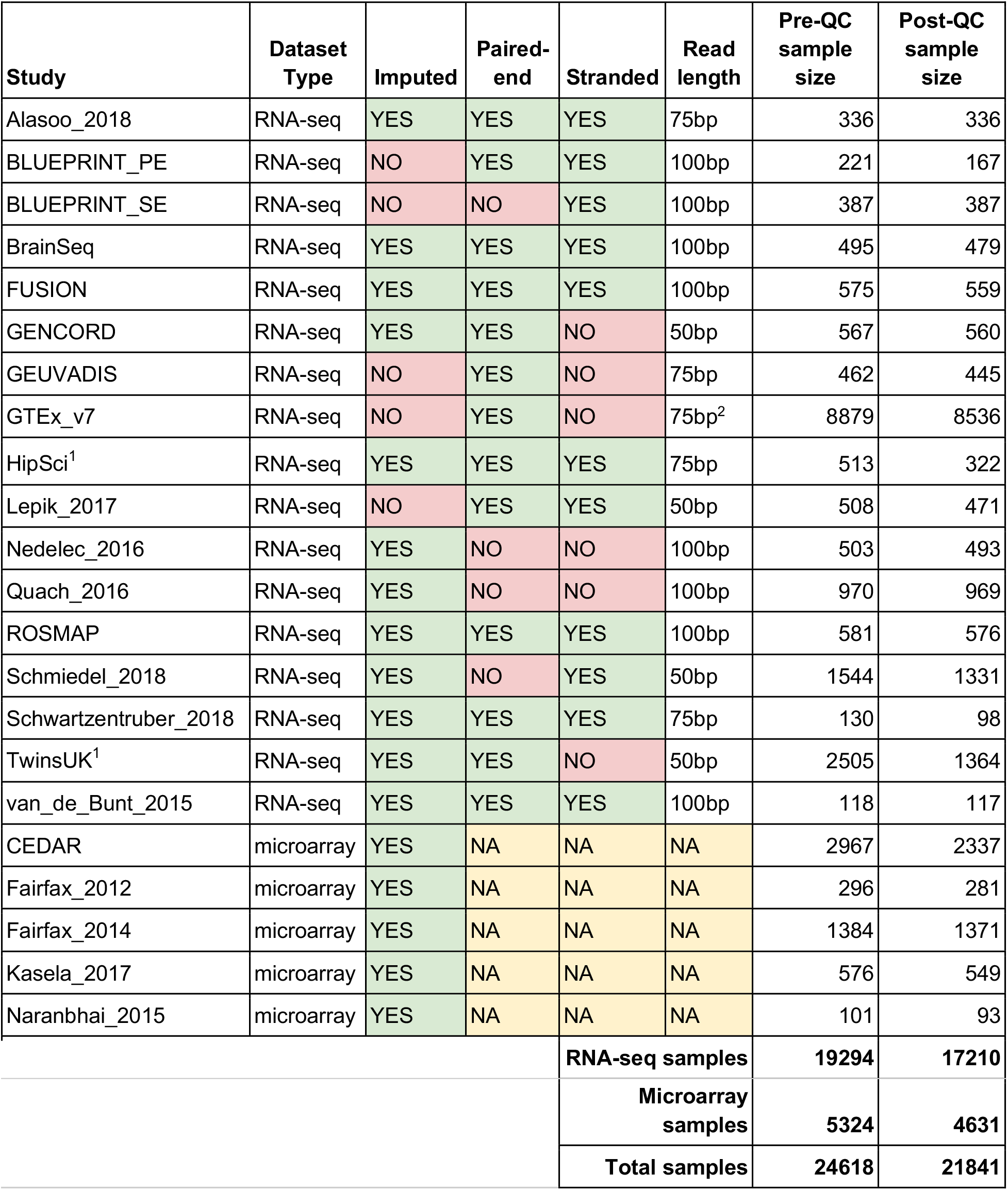
Overview of the studies included in the eQTL Catalogue. For four studies based on whole genome sequencing (BLUEPRINT, GTEx, GEUVADIS and Lepik_2017), we relied on final genotype files provided by the authors. All of the other genotypes were imputed using the 1000 Genomes Phase 3 reference panel (see Methods). ^1^TwinsUK and HipSci studies contain related individuals by design. These were excluded in the quality control step to enable eQTL analysis with a linear model. ^2^Small fraction GTEx samples have RNA-seq read lengths of 100 and 250 bp.

**Supplementary Figure 1.**
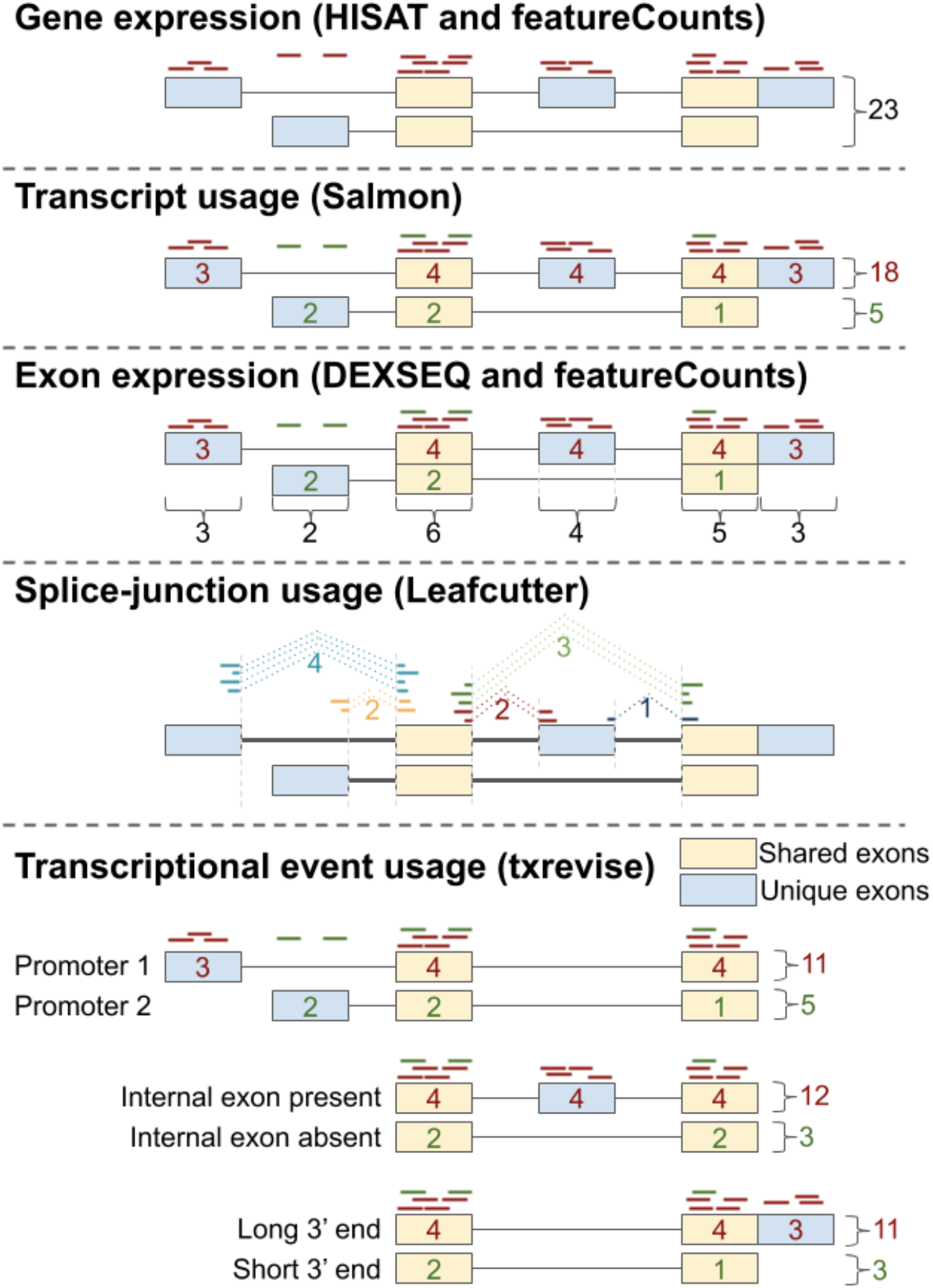
Quantification methods of molecular traits in the eQTL Catalogue. Symbolic representation of 23 read fragments assigned to 1 gene (aligned with HISAT2 *(72)*, quantified with featureCounts *(74)*) consisting of 2 transcripts (quantified with Salmon *(76)*) and 6 exonic parts (annotated with DEXSeq *(75)*, quantified with featureCounts). The gene also has 5 distinct introns which are identified and quantified by Leafcutter *(104)*. Transcriptional event usage is quantified with txrevise *(38)*. Txrevise uses shared exons as a scaffold to identify independent transcriptional events corresponding to alternative promoters, internal exons and 3’ ends. Leafcutter splice junction QTLs will be included in a future version of the eQTL Catalogue.

**Supplementary Figure 2.**
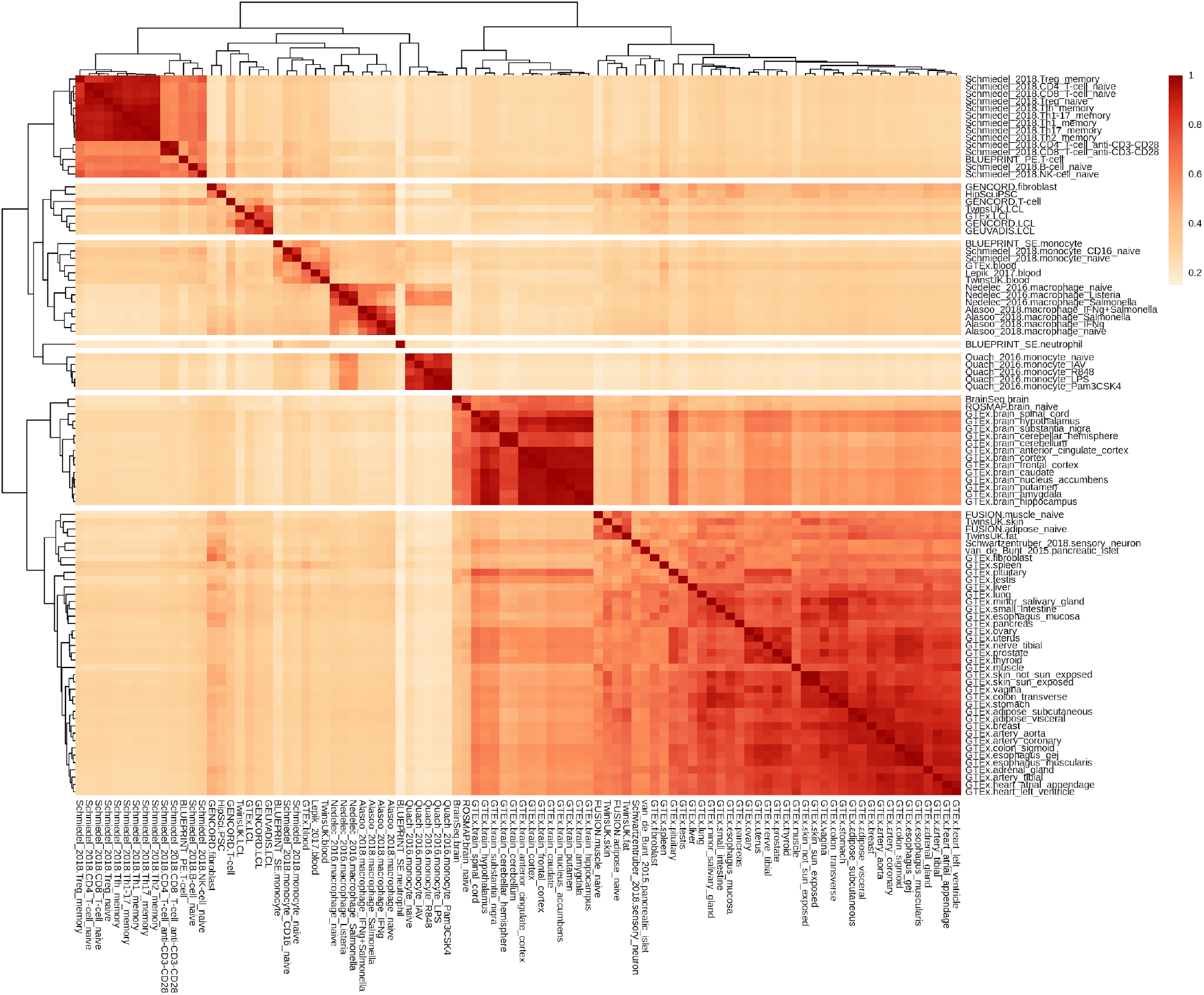
Pairwise eQTL sharing between 95 datasets estimated with the Mash model. We used 54,733 independent gene variant pairs from the fine mapping analysis (see Methods) and used the Mash model to estimate eQTL sharing between all pairs of the 95 datasets measured with RNA-seq.

**Supplementary Figure 3.**
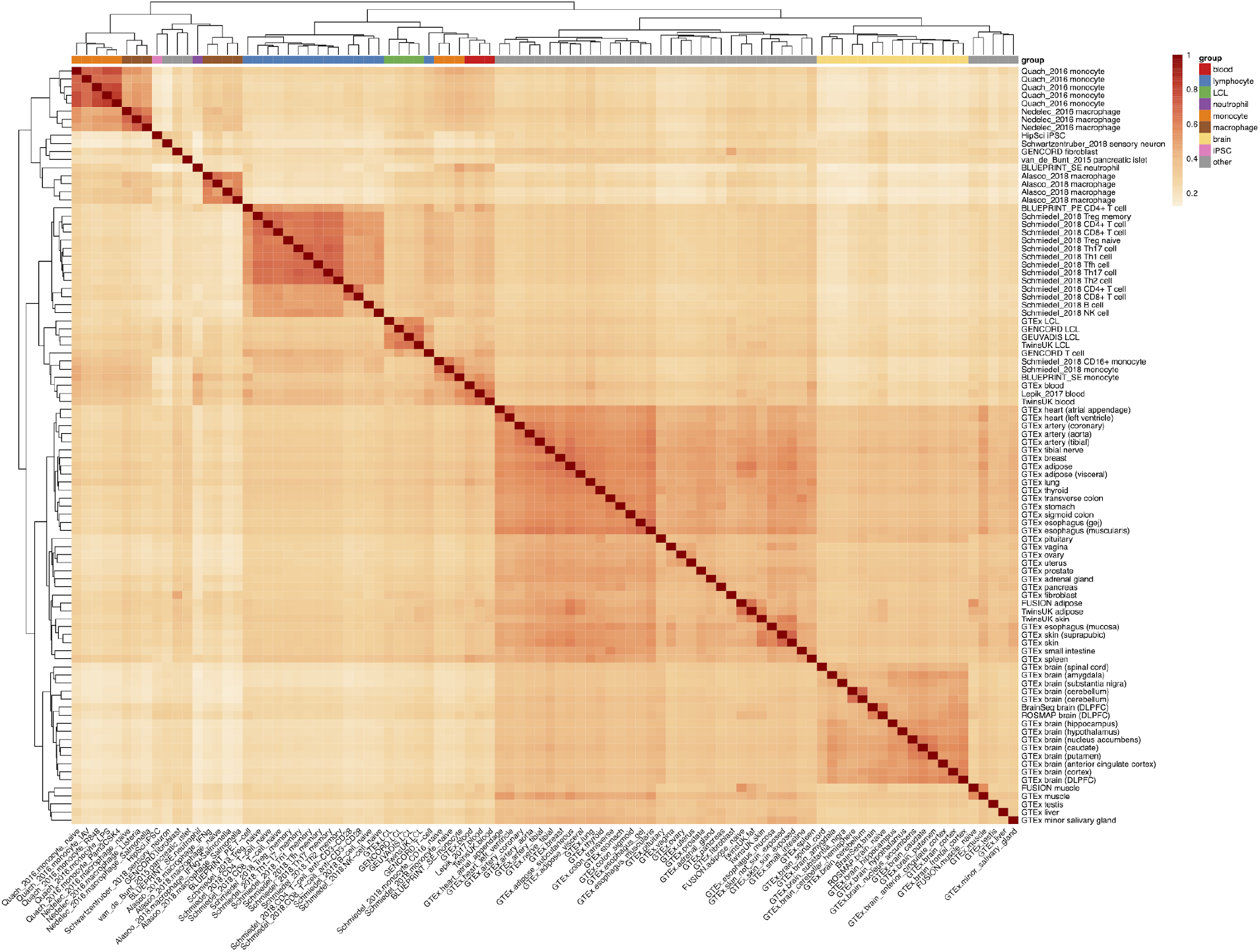
Pairwise eQTL similarity between 95 datasets estimated with Spearman correlation. We used 54,733 independent gene variant pairs from the fine mapping analysis (see Methods) and used the Spearman correlation of eQTL effect sizes to estimate eQTL sharing between all pairs of the 95 datasets measured with RNA-seq.

**Supplementary Figure 4.**
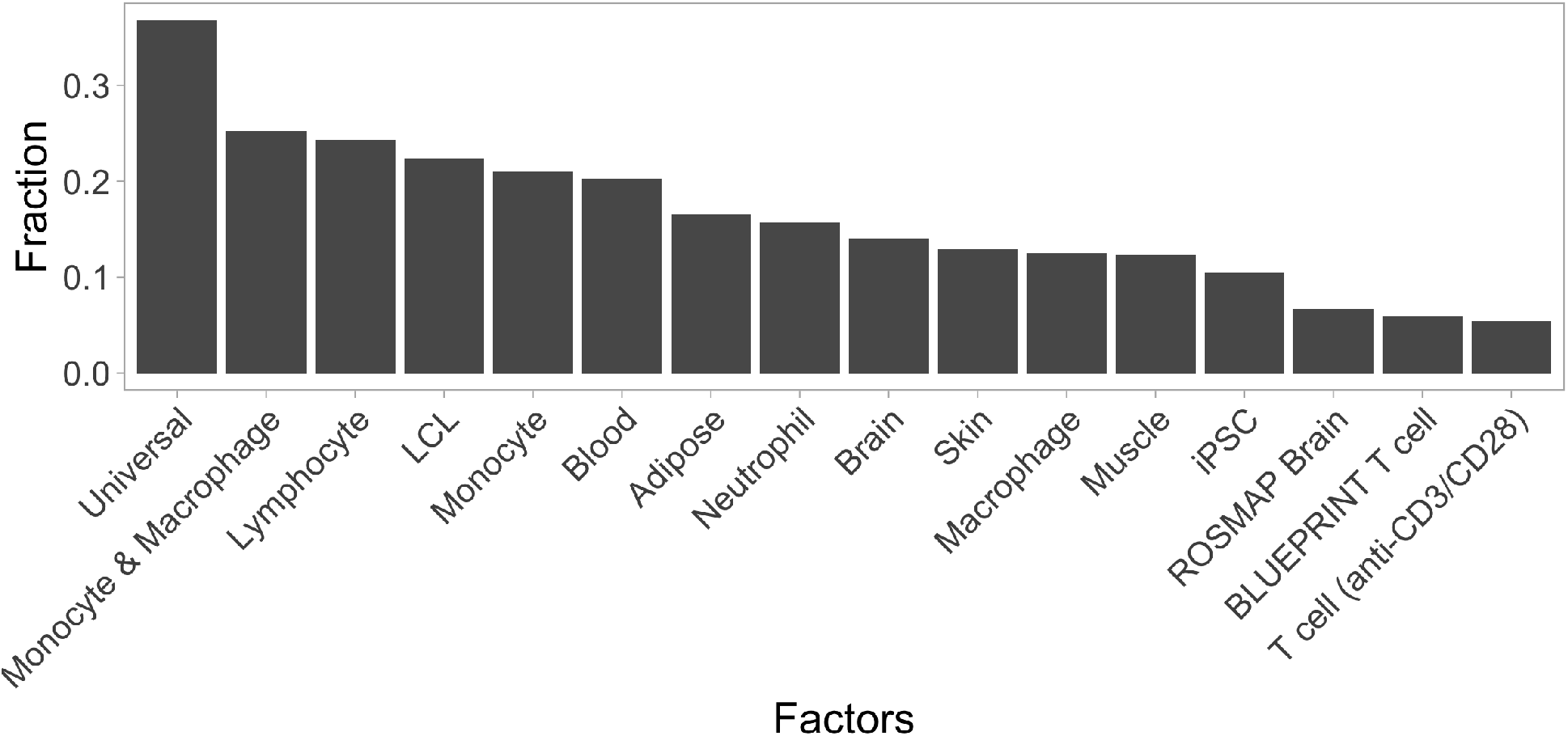
The fraction of fine mapped eQTLs assigned to each of the 16 factors detected by the sn-spMF method.

**Supplementary Figure 5.**
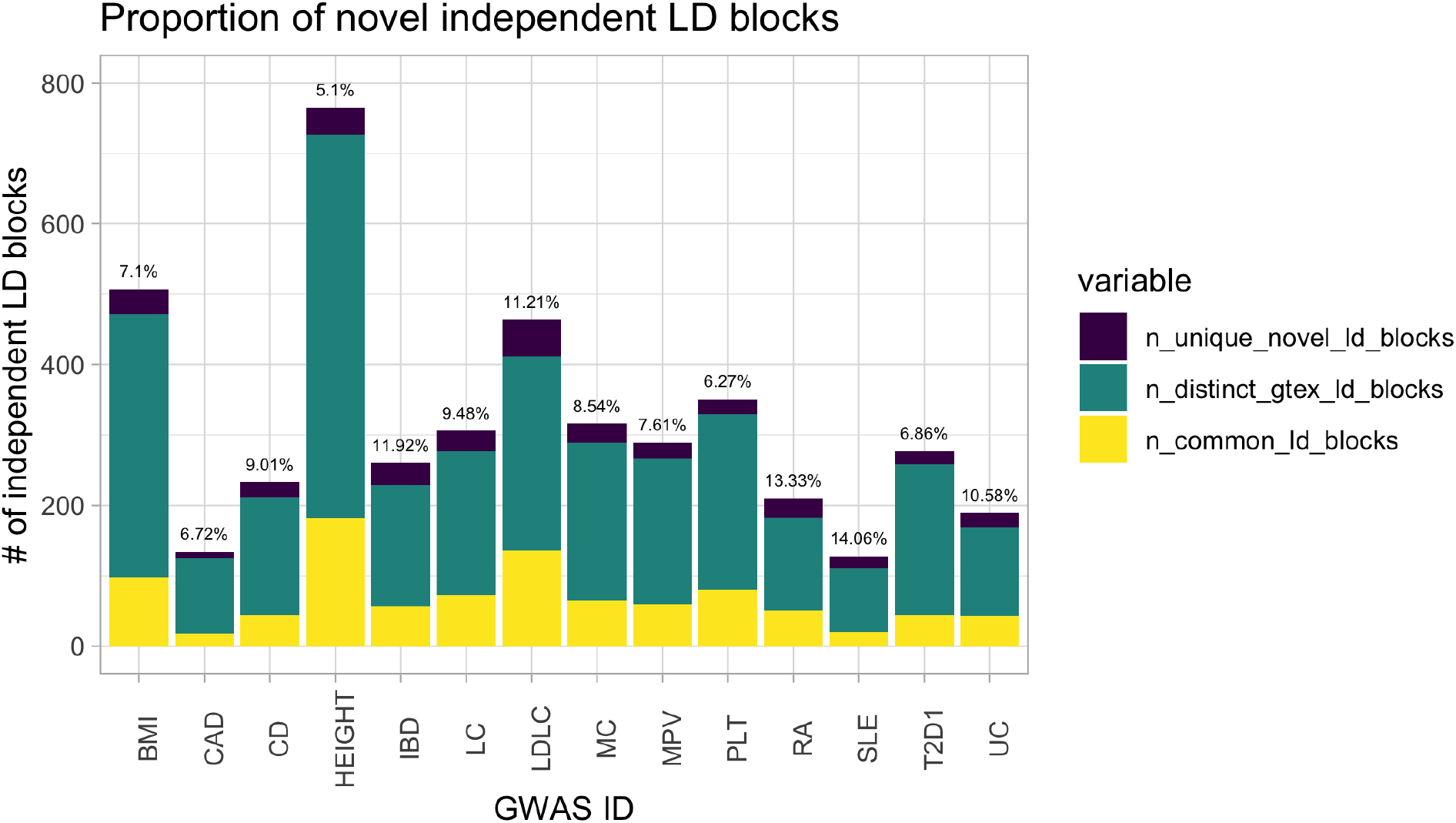
The number of shared and novel colocalisations detected for the 14 traits and diseases.

**Supplementary Figure 6.**
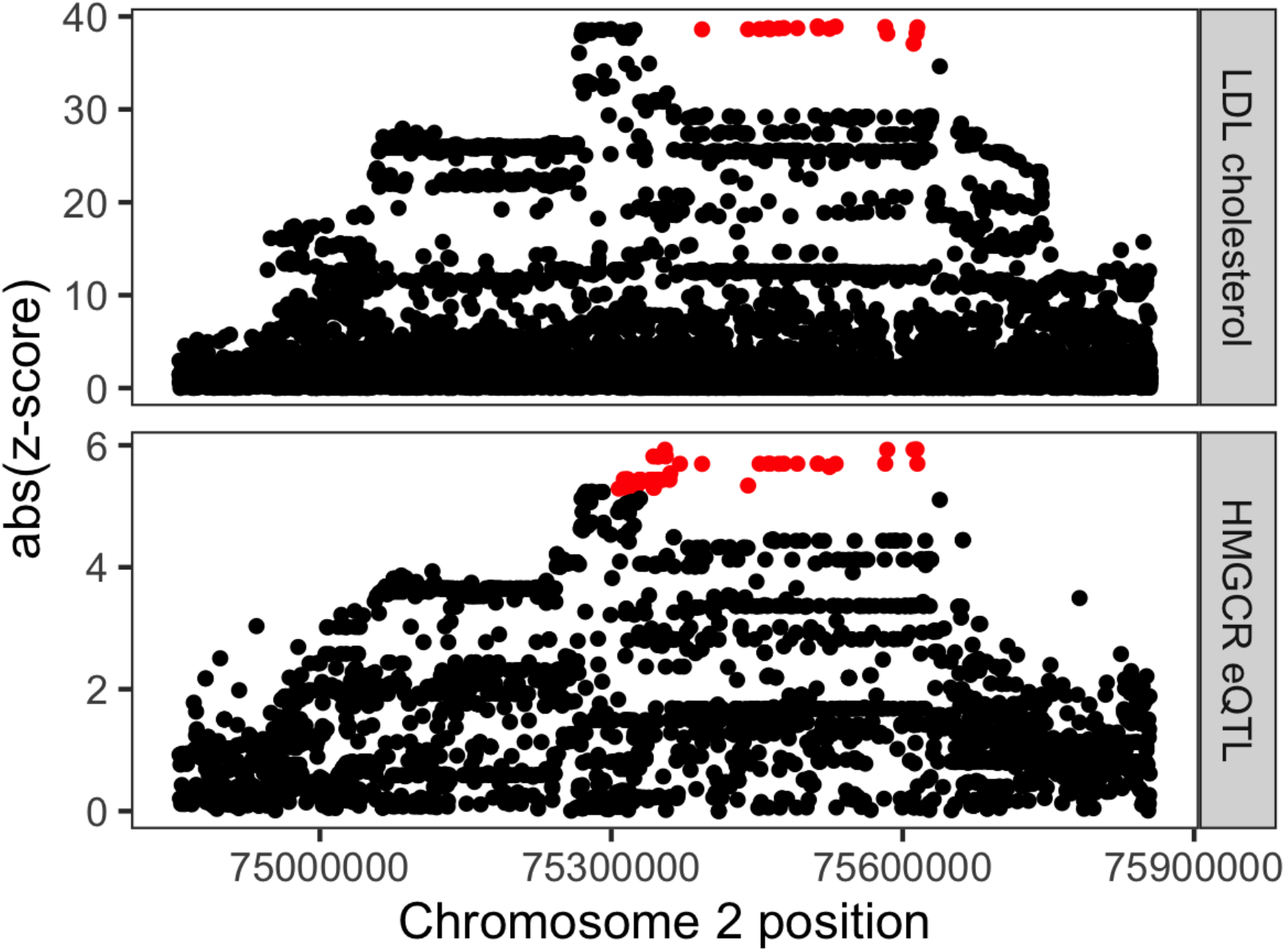
Regional association plot for LDL cholesterol (top panel) and *HMGCR* eQTL in the FUSION muscle dataset (bottom panel). The eQTL signal was fine mapped to 46 variants represented by red dots on both panels.

**Supplementary Figure 7.**
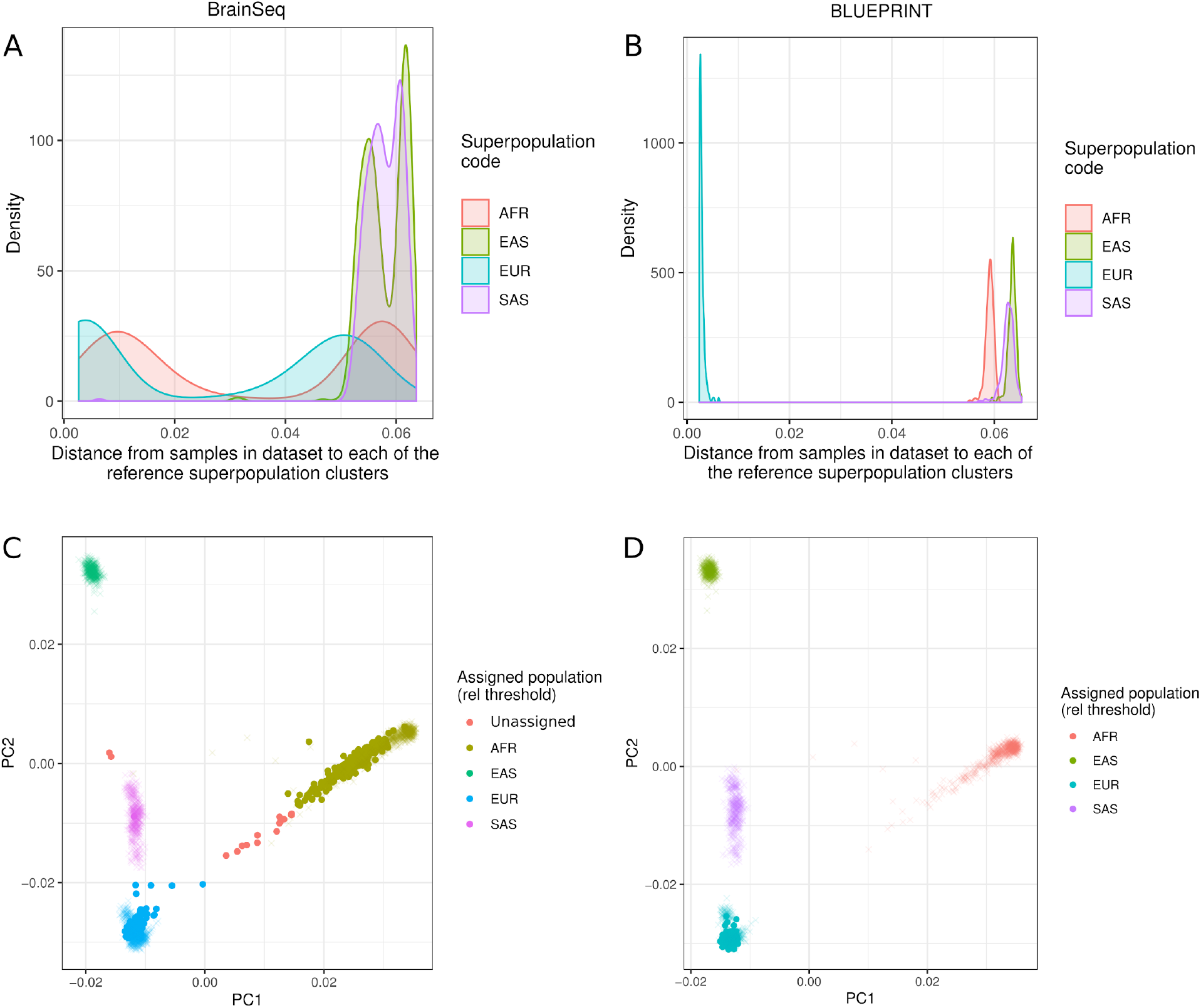
Assigning genotyped samples to the four 1000 Genomes superpopulations. **(A)**Density plot of distances between each sample in BrainSeq *(60)* dataset and each superpopulation cluster in the 1000 Genomes Phase 3 reference dataset *(30)*. First three principal components of the genotype data are used to calculate distances. The majority of samples in the BrainSeq dataset are close to either European (EUR) or African (AFR) superpopulations. **(B)**Histogram of distances between each sample in the BLUEPRINT *(20)* dataset and each superpopulation cluster in the reference dataset. All samples are close to the European (EUR) superpopulation cluster of the 1000 Genomes reference dataset. **(C)**Projection of the BrainSeq dataset to the first two principal components of the 1000 Genomes Phase 3 reference dataset. Most samples are assigned to either European or African superpopulations. Red samples are too far from all four superpopulations and thus remain unassigned. These samples are likely to represent recent admixture. **(D)**Projection of the BLUEPRINT dataset to the first two principal components of the 1000 Genomes Phase 3 reference panel. All samples are assigned to the European superpopulation. Superpopulation codes: EUR - European, AFR - African, SAS - South Asian, EAS - East Asian.

**Supplementary Figure 8.**
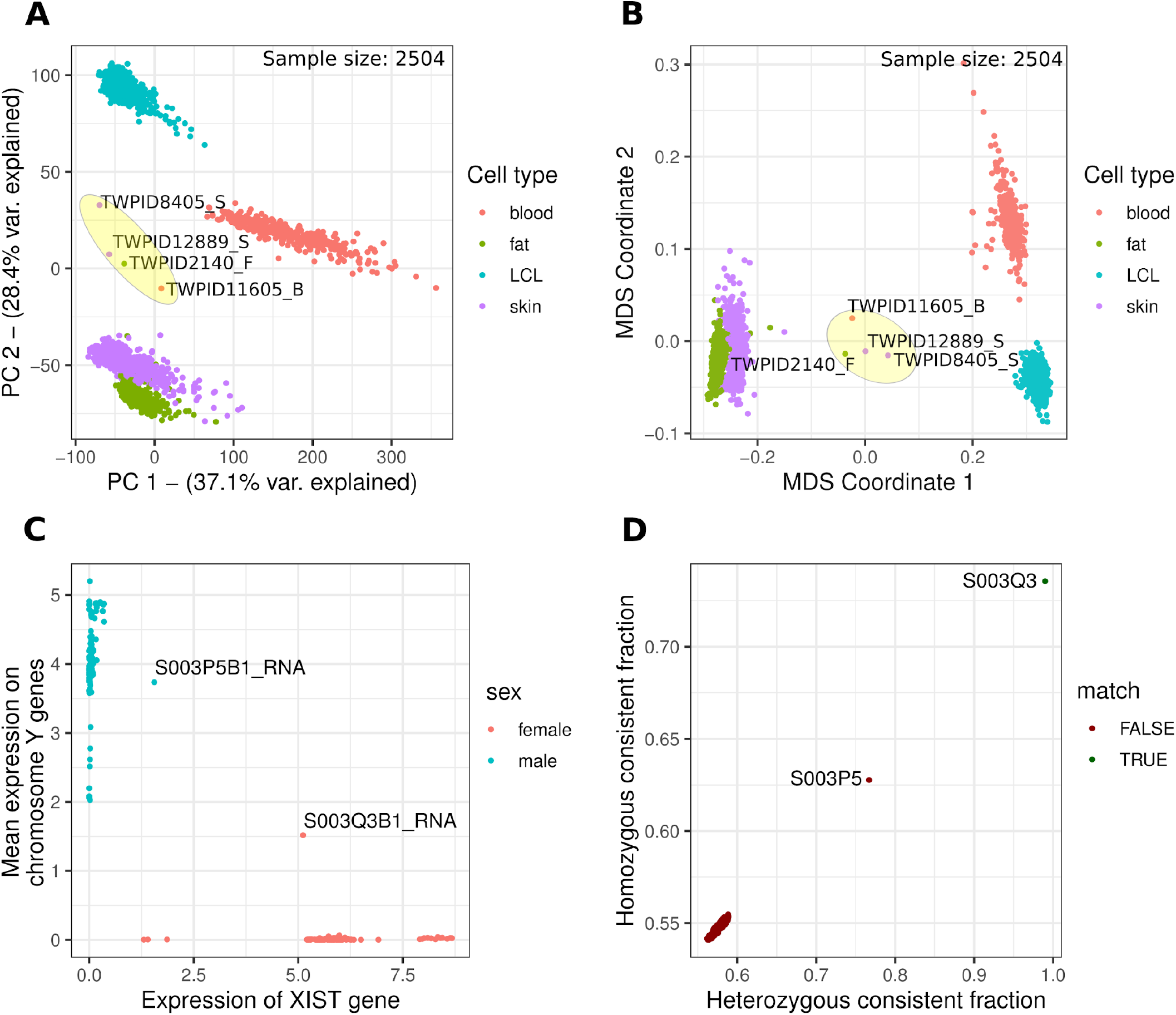
Overview of the Quality Control (QC) measures applied to all of the datasets in the eQTL Catalogue. QC reports for individual datasets can be found on the eQTL Catalogue website (https://www.ebi.ac.uk/eqtl/Studies/). **(A)** Principal component analysis of the TwinsUK dataset. **(B)** Multidimensional scaling analysis of the TwinsUK dataset. Four outlier samples (highlighted in yellow) from the PCA and MDS analysis were excluded from QTL mapping. (**C)** Sex-specific gene expression analysis. Expression of the female-specific *XIST* gene is plotted against the mean expression of the protein-coding genes on the Y chromosome. Samples from two donors (S003P5 (male) and S003Q3 (female)) expressed both *XIST* and genes from the Y chromosome, indicating potential cross-contamination with RNA from a sample of the opposite genetic sex. **(D)** Genetic similarity of S003Q3B1 RNA sample to all of the genotyped donors in the BLUEPRINT VCF file as calculated by the QTLtools mbv command *(81).* As expected, the genotypes of the S003Q3B1 RNA sample are most similar to the genotype data from the same donor (S003Q3) and most other donors are equally dis-similar, forming a separate cluster in the bottom left corner. However, the S003Q3B1 RNA sample also displays higher-than-expected genetic similarity with genotype data from the S003P5 donor. Together with the evidence presented in panel C, this suggests that cross-contamination has occurred between the S003Q3B1 and S003P5B1 RNA samples. As a result, we decided to remove these two samples from downstream analysis.

